# Weak and uneven associations of home, neighborhood and school environments with stress hormone output across multiple time scales

**DOI:** 10.1101/705996

**Authors:** Margherita Malanchini, Laura E. Engelhardt, Laurel Raffington, Aditi Sabhlok, Andrew D. Grotzinger, Daniel A. Briley, James W. Madole, Samantha M. Freis, Megan W. Patterson, K. Paige Harden, Elliot M. Tucker-Drob

## Abstract

The progression of lifelong trajectories of socioeconomic inequalities in health and mortality begins in childhood. Dysregulation in cortisol, a stress hormone that is the primary output of the hypothalamus-pituitary-adrenal (HPA) axis, has been hypothesized to be a mechanism for how early environmental adversity compromises health. However, despite the popularity of cortisol as a biomarker for stress and adversity, little is known about whether cortisol output differs in children being raised in socioeconomically disadvantaged environments. Here, we show that there are few differences between advantaged and disadvantaged children in their cortisol output. In 8- to 14-year-old children from the population-based Texas Twin Project, we measured cortisol output at three different time-scales: (1) diurnal fluctuation in salivary cortisol (*n* = 400), (2) salivary cortisol reactivity and recovery after exposure to the Trier Social Stress Test (*n* = 444), and (3) and cortisol concentration in hair (*n* = 1,210). These measures converged on two moderately correlated, yet distinguishable, dimensions of HPA function. We then tested differences in cortisol output across nine aspects of social disadvantage at the home (*e.g.*, family socioeconomic status), school (*e.g.*, average levels of academic achievement), and neighborhood (*e.g.*, concentrated poverty). Children living in neighborhoods with higher concentrated poverty had higher diurnal cortisol output, as measured in saliva; otherwise, child cortisol output was unrelated to any other aspect of social disadvantage. Overall, we find limited support for alteration in HPA axis functioning as a general mechanism for the health consequences of socioeconomic inequality in childhood.

## INTRODUCTION

As income inequality in the United States widens, disparities in health and survival between people in the bottom versus the top of the socioeconomic distribution continue growing.^1–4^ Motivated by the goal of understanding, and ultimately mitigating, socioeconomic disparities in health outcomes, the biosocial research agenda has attempted to identify specific biological mechanisms for how exposure to disadvantage gets ‘under the skin’^5^ to produce sub-optimal life outcomes. In this effort, perhaps no other biomarker has been more widely studied than cortisol.^6^ Cortisol is the human glucocorticoid that is the major output of the hypothalamus-pituitary-adrenal (HPA) axis of the neuroendocrine system, which regulates a suite of physiological processes, including immune function, metabolism, cardiovascular function, and central nervous system function, and is highly responsive to both psychological and physical stress.^7^ Given the extensive investment of scientific resources into cortisol research, and the gaping health inequalities between poor and rich Americans,^3^ it is essential for researchers to be able to make informed choices about which measures of cortisol output are most robustly associated with socioeconomic inequalities.

The hypothesis that glucocorticoid response is a critical mechanism for the biological embedding of stress^8–10^ is grounded in over six decades of animal research^11^ demonstrating that early environmental exposure changes the HPA response to stress.^12,13^ For instance, adult rats exposed to periods of stimulation^14^ and maternal care^15,16^ during the first few weeks of life exhibit reduced glucocorticoid responses to stress compared with non-stimulated animals. Two lines of evidence in human studies further support the hypothesis that glucocorticoid response is a translational mechanism for the biological embedding of stress. First, cortisol output has been associated with mental and physical health, including sleep disturbances, depression, obesity, and cardiovascular disease.^17–19^ Second, basal salivary cortisol has been linked to a range of chronic or severe psychological stressors, most notably neglect, abuse and maltreatment in childhood.^20–23^

Motivated by these findings, cortisol measurement has been incorporated into large-scale epidemiological research aimed at elucidating biological markers of health and pre-disease state. Notably, however, few of these studies have reported associations between cortisol output and measures of socioeconomic disadvantage. In their review of large-scale epidemiological investigations of cortisol, Adam and Kumari (2009) identified only 2 such studies (out 17 total), both of which focused on cortisol diurnal rhythm measured in adults on a single day.^24^ One study found that adults with lower income and educational attainment had higher diurnal cortisol output.^25^ Similarly, the second found that older adults with lower occupational status and wealth had higher diurnal cortisol output.^26^ More recently, another study also found that adults with lower socioeconomic status show flatter diurnal rhythm, characterized by lower peak after awakening and higher levels of cortisol in the evening.^27^

However, other research linking cortisol output to socioeconomic disadvantage has found inconsistent results. Some studies have failed to find expected associations between disadvantage and higher cortisol output (^22^ for a review) and still others found that socioeconomic adversity was associated with *hypo*cortisolism rather than *hyper*cortisolism.^28,29^ Mixed evidence of an association between socioeconomic deprivation and diurnal cortisol rhythm also comes from a series of observational^30^ and experimental studies^31^ of adult samples in Kenya, and the generalizability of these results to understanding social inequalities in Western samples is unclear. Some theories have attempted to reconcile these inconsistences by proposing non-linearity in the association between the severity of socioeconomic disadvantage and cortisol output.^32,33^

Evidence for an association between cortisol output and socioeconomic disadvantage is even scarcer for children, as the majority of studies in child samples have focused instead on extreme forms of psychological and/or physical stress – most commonly, neglect and maltreatment. While research in this area has been advanced by two meta-analyses,^23,34^ these meta-analyses provide little insight into how socioeconomic disadvantage, specifically, is linked to variation in cortisol output. Moreover, the meta-analyses indicated that publication bias in the childhood hormonal literature was likely, which could have led to inflated estimates of effect size and significance.^23,34^ One of the most influential studies supporting the association between cortisol and socioeconomic status was published nearly two decades ago.^5^ In 6- to 10-year-old children, but not 12- to 16-year old children, family income, education and employment were associated with higher morning salivary cortisol. More recently, an association between blunted reactivity to stressors and lower family income was observed in a sample of 6-7 year-olds.^35^ The dearth of studies focused on socioeconomic advantage in childhood is problematic, as socioeconomic inequalities in adult health and mortality are rooted in childhood experiences.^36^

Efforts to understand how child cortisol output is related to socioeconomic disadvantage are further stymied by the fact that cortisol output can be measured in multiple ways. First, cortisol reactivity/recovery is the ability of the HPA axis to produce an adaptive response when exposed to an acute stressor.^37^ Normative reactivity/recovery profiles are characterized by substantial increases in cortisol in response to an environmental stressor, accompanied by a rapid rate of recovery back to baseline when exposure has ended, typically within 40-60 minutes.^38^ High reactivity, slower than expected recovery, or an overall blunted reactivity profile have all been considered maladaptive. Second, daily cortisol production is marked by a pronounced diurnal rhythm, with levels rising through the night, peaking between 30-45 minutes after waking and then declining over the rest of the day.^39,40^ Slower rates of cortisol decline throughout the day, which correspond to higher evening levels, are considered maladaptive.

Finally, researchers can measured trait-like differences in overall levels of cortisol, with both unusually high^41,42^ and unusually low^43^ basal levels linked to maladaptive health outcomes. Stable individual differences in cortisol concentration can be measured by aggregating across multiple repeated salivary or urinary samples collected at different times in the day (*e.g.*,^44^), or by measuring cortisol concentration in hair.^45^ Heightened cortisol levels, thought to reflect protracted stress exposure, have been hypothesized to be more strongly associated with environmental adversity than other cortisol indices.^19,46^ However, a systematic review of 15 studies, with a median sample size of 242, found that the evidence for a link between hair cortisol concentration and socioeconomic disadvantage was only suggestive.^47^

Despite the availability of multiple measures of cortisol output, previous research has commonly collected, reported and/or interpreted results for only a single measure.^11,48^ But, it is doubtful that different measures can be treated as interchangeable indicators of the same underlying aspect of HPA functioning.^44^ One study of just 17 adults found little correspondence between cortisol concentration in hair and daily fluctuation in salivary cortisol.^44^ Another study found that shallower decreases in diurnal slope were modestly associated with lower reactivity to and longer recovery after the Trier Social Stress Test (TSST).^49^ A few other extant studies all included fewer than 35 people.^50,51,52^

There has not yet been a comprehensive effort to measure multiple dimensions of cortisol output *and* socioeconomic disadvantage in a single, well-powered sample of children. This approach is necessary to identify which aspects of HPA axis functioning are most strongly associated with which dimensions of child socioeconomic disadvantage, while accounting for the overlap between constructs and correcting for multiple testing and researcher degrees of freedom. In the present study, we use 17-20 cortisol samples per child to examine 7 reliable indices of cortisol output across 3 time-scales. Furthermore, we consider multiple indices of the home, school, and neighborhood environment.^53^ This comprehensive approach provides an important test of a popular paradigm for research that aims to understand the biological mechanisms for how environmental adversity gets ‘under the skin.’ Progress on this research goal is particularly important, given the widening gap in physical, psychological and psychiatric health between the high and low ends of the socio-economic distribution.^1–3,10,54^

## RESULTS

We studied the relationship between socioeconomic disadvantage and cortisol output in 8- to 14-year-old children from the Texas Twin Project who were recruited from public school rosters (*n* = 400 - 1,210 depending on cortisol metric). Figure 1 summarizes the measures of cortisol and social context, as well as the analytic approach. We tested the associations between socioeconomic disadvantage and cortisol output at three levels of granularity, moving from broad constructs that reflect communalities across cortisol measures and all socioecological context measures, to specific associations between individual measures.

**Figure 1.**
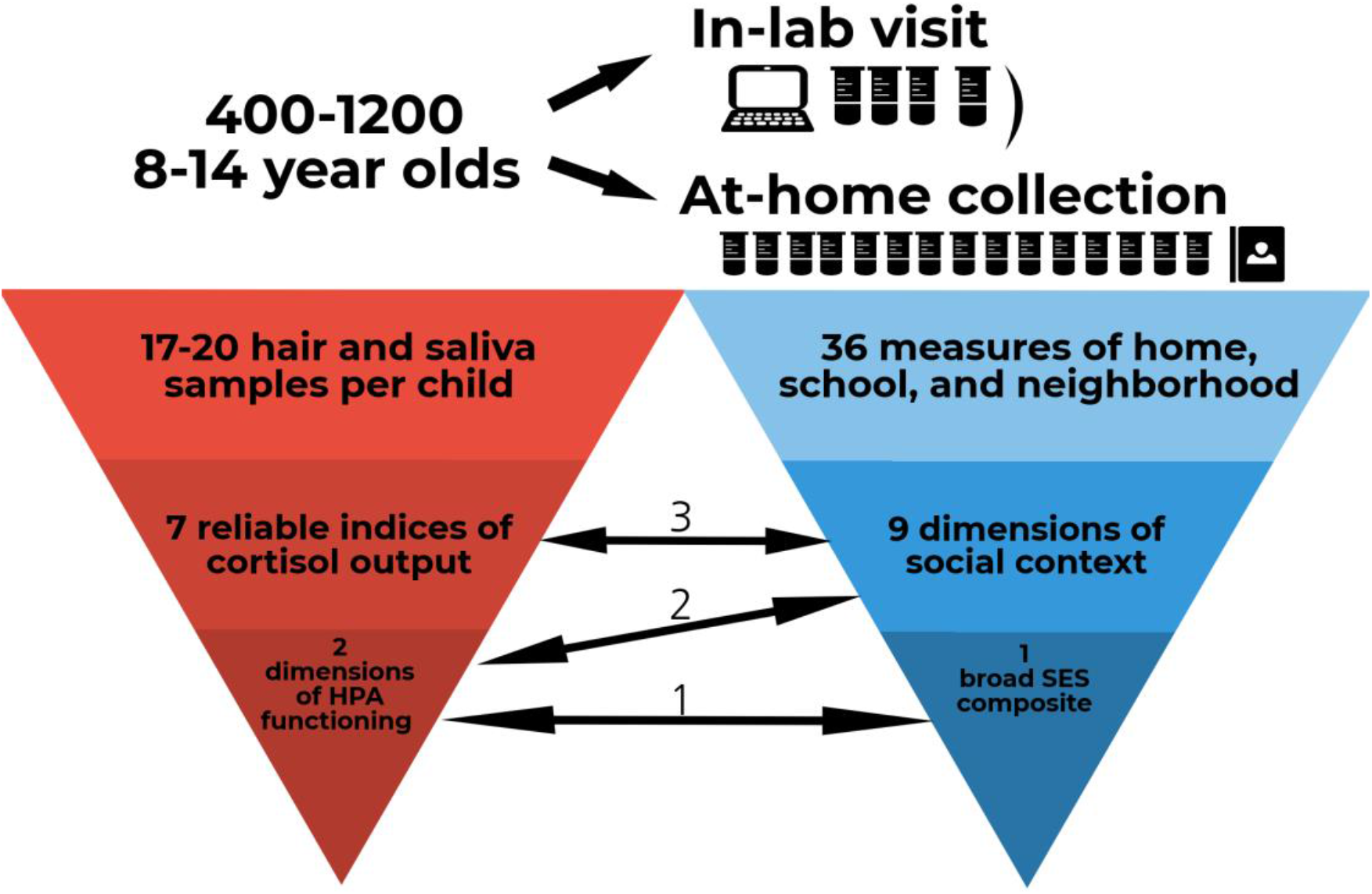
Details of the approach we adopted for measuring variation in cortisol and socioecological disadvantage.

Although all children were recruited from a single metropolitan area, inequalities in children’s social contexts were stark. The Gini index (calculated using the *gini* function of the R package ‘DescTools’^55^) of the income distribution in this sample was 0.35. This estimate is very similar to the Gini coefficient for the United States as a whole in 2016 (0.39), albeit slightly lower- as would be expected for a regional compared to national sample. This coefficient is also comparable to coefficients for other developed countries including Israel, Latvia and New Zeeland.^56^ We previously reported that information from parent reports, state educational agency data, and U.S. census data can be integrated into 9 dimensions of social context (^53^; Methods). At the neighborhood level, children varied in their exposure to *concentrated poverty*, *residential instability*, and *race/ethnic diversity*. At the school level, children varied in their exposure to *low academic achievement*, *teacher inexperience*, and *race/ethnic diversity*. At the home level, children varied in their *family socioeconomic status*, and their exposure to *cumulative adversity* (financial difficulties and life events) and *interparental conflict*. These measures of sociecological disadvantage were weakly to moderately correlated (Figure S1).

Children participated in a research laboratory visit that included a Trier Social Stress Test (TSST),^38^ which required children to prepare and present a short story and do some mental arithmetic in front of an unfamiliar audience comprising two “judges”. Salivary hormonal samples were taken before (1) and after (3) the TSST. Children also contributed a hair sample during the lab visit and then completed a hormonal sampling protocol at home, requiring them to contribute 3 salivary samples a day for 4-5 days. This resulted in a total of 17-20 saliva and hair samples per child. From these samples we extracted 7 reliable indices of cortisol output reflecting variation in diurnal cortisol rhythm, acute stress reactivity and recovery, and trait-like hair cortisol concentration.

### Repeated measures of cortisol capture two distinct dimensions of HPA functioning

We used multi-level piecewise growth models to analyze repeated salivary measures from home sampling and the lab visit, in order to model individual differences in diurnal rhythm and in acute stress response, respectively. Compared to previously-used analytic approaches, such as area under the curve (AUC,^57,58^ and traditional latent growth models^49,59^), this approach accurately captures individual variation in rates of change throughout the day and throughout the experience of an acute stressor, while taking into account timing and spacing of repeated sampling (see Methods and Supplementary Information).

Results of the multi-level piecewise growth models indicated that children varied substantially in their cortisol trajectories across days at home (Figure 2a) and across minutes in the laboratory (Figure 2b). Analyses of diurnal change found that children showing greater cortisol awakening responses had lower cortisol levels at waking (*r* = −.61, *p_uncorrected_* = .049) and maintained higher levels of cortisol throughout the day (*r* = .49, *p_uncorrected_* = .015). Analyses of change in response to acute social stress found that children with higher levels of cortisol before the TSST also showed less reactivity to the TSST-induced stress (*r* = −0.43, *p_uncorrected_* = .0005) but did not differ in the rate of their subsequent recovery in cortisol post stressor (*r* = −0.05, *p_uncorrected_* = .515). Children with heightened cortisol reactivity showed faster cortisol recovery (*r* = −0.57, *p_uncorrected_* = .0005).

**Figure 2.**
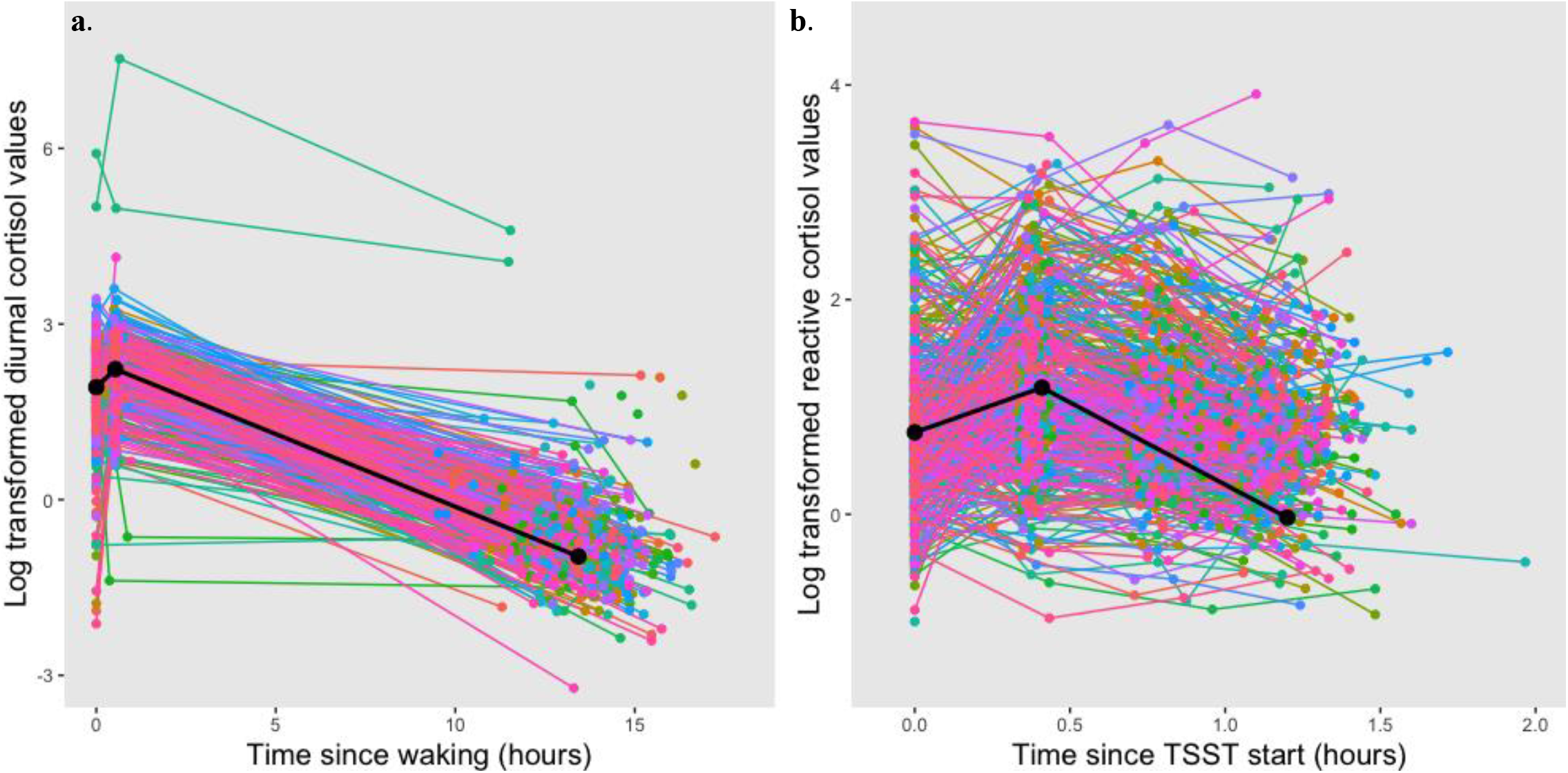
Patterns of within and between individual variability in diurnal and reactive cortisol trajectories calculated applying multi-level piecewise growth models. Figure 2a shows the individual trajectories (colored lines) and the mean trajectory (black line) of diurnal variation in cortisol, based on the first day of sampling. The same plots for the five consecutive days are reported in supplementary Figure S2. Figure 2b shows the individual trajectories (colored lines) and mean trajectory (black line) of cortisol output over the period of time surrounding the Trier Social Stress Test (TSST). All estimates are calculated after applying the exclusion criteria and accounting for potentially confounding covariates described in the supplementary information, and including age, sex and age×sex as between-level correlates to the model. Full model results can be found in supplementary tables S1 and S2.

We then assessed the extent to which diurnal cortisol output covaried with cortisol response to acute stress, as well as with cortisol concentrations measured in hair. Correlations among the 7 indices of cortisol output are presented in Figure 3 and Supplementary Table S3. Children with higher levels of hair cortisol showed flatter diurnal slopes from morning to evening (*r* = 0.27, *p_uncorrected_* = .0001), higher pre-TSST levels in the lab (*r* = .31, *p_uncorrected_* = .020), and slower TSST reactivity (*r* = −.16, *p_uncorrected_* = .008). Children with flatter diurnal slopes from morning to evening (*i.e.*, maintaining higher levels of cortisol throughout the day) also showed higher pre-TSST cortisol levels in lab (*r* = 0.22, *p_uncorrected_* = .004). Most associations remained significant after accounting of multiple testing using the Bejamini-Hochberg false discovery rate (FDR) method^60^, calculated using the *p.adjust* function in R (see Table S3), with the exception of the correlations between awakening response and diurnal slope (*r* = .49, *p_corrected_* = .059) and between hair cortisol and pre-TSST levels (*r* = .31, *p_corrected_* = 0.060). Overall, indicators of trait-like elevations in cortisol levels (hair, diurnal slope, and pre-TSST cortisol output) were modestly inter-correlated but showed little correspondence with cortisol responsiveness either to an acute stress or to waking

**Figure 3.**
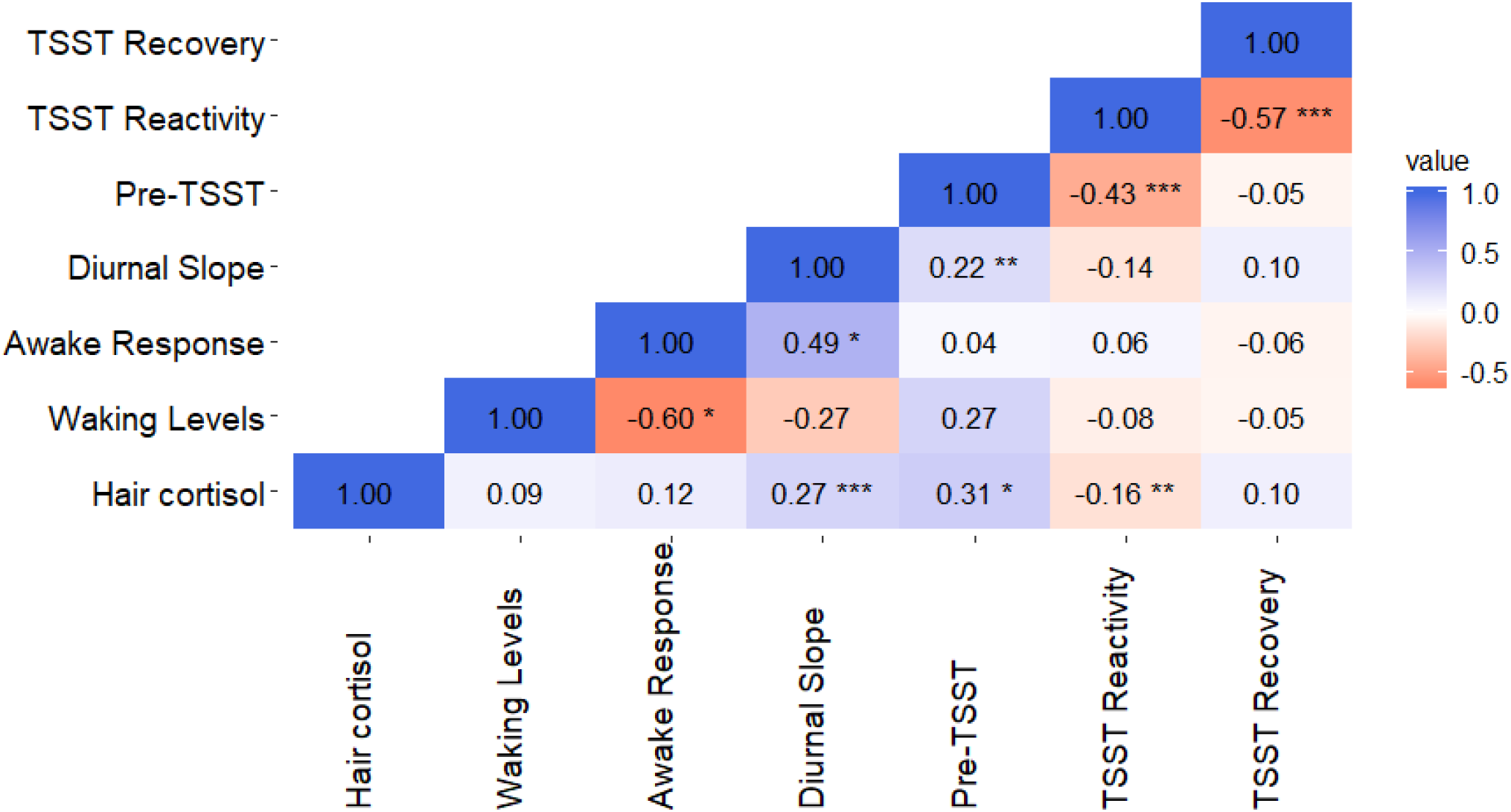
Correlations between variation in diurnal and reactive cortisol rhythm and hair cortisol. We simultaneously modelled diurnal and reactivity/recovery trajectories using multivariate growth modelling, as a tool examine their correspondence. All associations were calculated after controlling for potentially confounding covariates (including batch, medication use, eating and drinking and dairy intake, see supplementary information for a detailed account), and including age, sex, age×sex and race, as correlates to the model; * = *p*< .05, ** = *p*< .01, *** = *p*< .001.

Based on the correlations among all cortisol indicators (Figure 3), we fitted a second-order latent factor model with two dimensions of HPA functioning: A *pervasive* and a *diurnal* factor of cortisol output (Figure 4 and Table S4). Loading on the *pervasive* factor were higher hair cortisol (*λ* = .474, *p* < .001), higher pre-TSST cortisol levels (*λ* = .663, *p* < .001) and flatter diurnal slope from morning to evening (*λ* = .699, *p* < .001). Variation in cortisol reactivity to TSST also had a small, negative loading on the *pervasive* factor (*λ* = −.236, *p* < .001). This *pervasive* factor of HPA function was interpreted to reflect stable individual differences in HPA function, potentially indexing a biological signature of long-term stress exposure manifesting as steadily heightened levels of cortisol across multiple assays.

**Figure 4.**
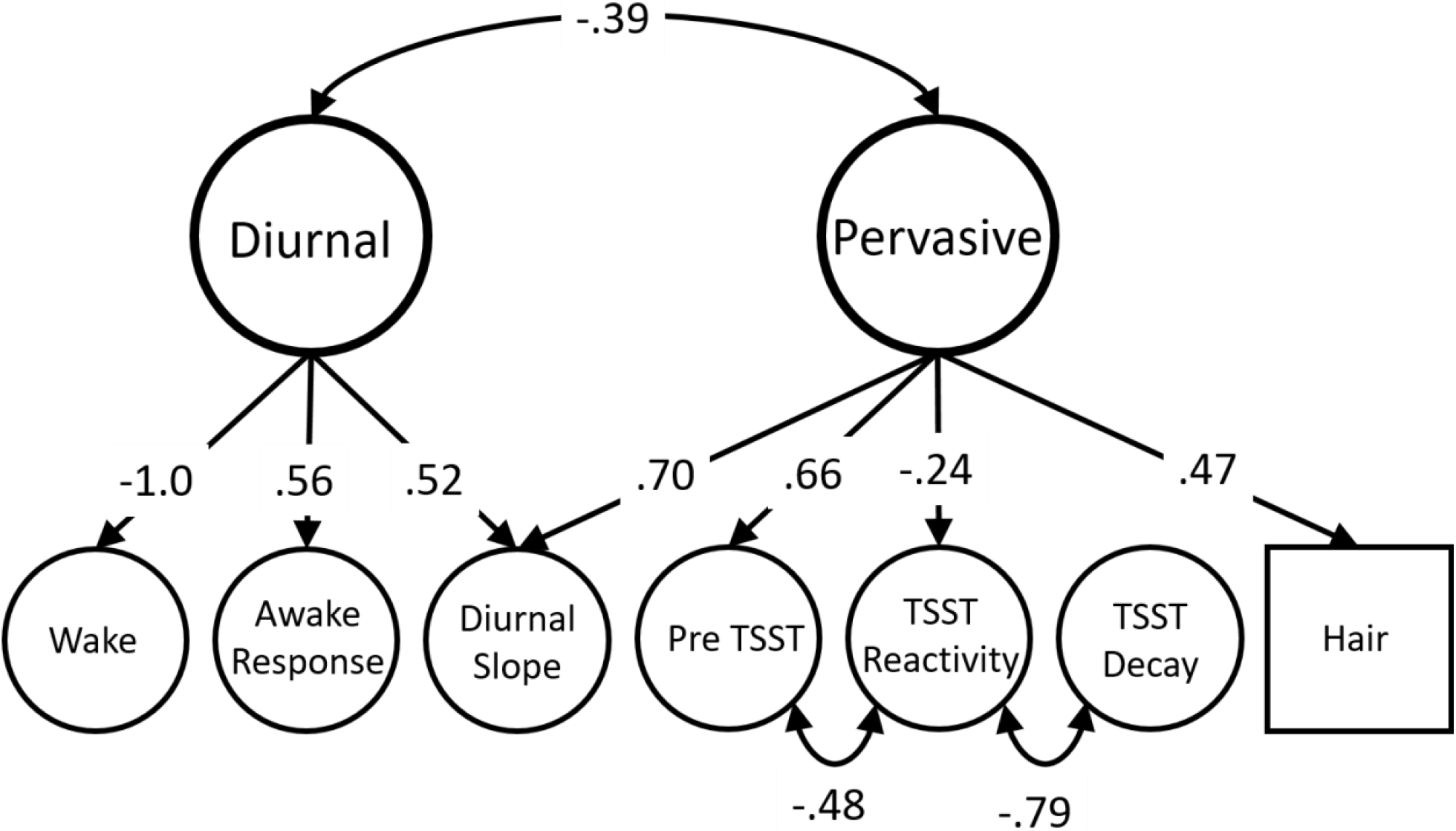
Association between second order latent factors of cortisol, clustering variation common across latent growth parameters into chronic and diurnal cortisol levels, all paths are standardized, * = p< .05, ** = p< .01. The two latent factors of HPA functioning are second order latent factors, which were obtained fitting a hierarchical model on top of the two multi-level piecewise growth models for diurnal and reactive cortisol with hair cortisol added as a between-level correlate (see Methods). Therefore, these two latent dimensions of HPA function represent the variance common to the seven reliable indices of cortisol output (six latent indices obtained from growth models and one observed index of hair cortisol), which were themselves obtained from the 17-20 cortisol samples collected at home and in-laboratory.

The *diurnal* factor was interpreted to reflect variation in naturally-occurring diurnal fluctuation of cortisol. Loading onto the *diurnal* factor were the three growth parameters indexing variation in diurnal rhythm: lower waking levels (*λ* = −.998, *p* < .001), higher awakening response (*λ* = .559, *p* < .001), and flatter diurnal slope (*λ* = .516, *p* < .01). Post-TSST recovery did not load on any latent cortisol factors. However, its strong negative association with reactivity after stress exposure (*r* = −.793, *p* < .001), and the negative correlation between pre-TSST levels and reactivity to the in-lab stressor beyond the common *pervasive* factor (*r* = −.475, *p* < .001) were captured with residual correlations (Figure 4). These two higher-order dimensions of HPA functioning were only moderately correlated (*r* = −.394, *p* < .05).

### The relationship between cortisol output and social context is specific to neighborhood concentrated poverty

We first estimated the association between social context and cortisol output at the broadest level: Do children who are generally disadvantaged across all environmental settings show differences in cortisol output? This analysis focused on the two higher-order factors of cortisol output (*pervasive* and *diurnal*). A single composite measure of general disadvantage was constructed by conducting a principal component analysis (PCA) of the 9 dimensions of social context and then calculating a mean score weighted by their loadings on the first principal component, which explained 30.0% of the variance. Considering variation in cortisol output and social context through this wide lens, neither the *diurnal* nor the *pervasive* factor was significantly related to general disadvantage (*r* = −.008, *p_uncorrected_* = .933, and *r* = −0.179, *p_uncorrected_* = .075, respectively).

Given the complexity of social context, the previous wide-lens analysis might obscure important associations between HPA function and specific aspects of disadvantage. Consequently, we next examined the associations between the two higher-order cortisol factors and each of the nine dimensions of social context (Table S5). Results showed that children living in advantaged neighborhoods with*out* concentrated poverty had lower *diurnal* cortisol (*β* = −.190, *p_uncorrected_* = .002) but did not differ in *pervasive* cortisol output. That is, children living in neighborhoods with higher concentrations of poverty showed lower waking levels of cortisol, more pronounced awakening responses, and flatter declines in cortisol over the course of the day. The association between neighborhood poverty and *diurnal* cortisol remained significant after correcting for multiple comparisons using the Bejamini-Hochberg FDR method (*β* = −.190, *p_corrected_* = 0.035). No other dimension of social context was associated with either HPA function factor.

As extant literature on the links between cortisol output and environmental adversity has suggested that these relations are characterized by non-linear associations (27, 28), we examined whether the pattern of relations changed when we added quadratic terms for the nine socioecological indicators to each model. School-level achievement showed a significant negative quadratic association with the pervasive factor of cortisol output (*p_uncorrected_* = .003, see supplementary Table S6), but not with the diurnal factor. Children attending schools characterized by moderate levels of achievement showed higher pervasive cortisol output, while children attending schools with either very high or very low levels of achievement showed diminished levels of pervasive cortisol output (see supplementary Figure S3). However, this association was not significant after we corrected for multiple testing using the Bejamini-Hochberg FDR method (*β* = −.356, *p_corrected_* = .0540). No other socioecological indicator was associated with either of the two higher-order cortisol dimensions.

Earlier research has argued for the existence of a sensitive period in which associations between cortisol and socioeconomic status are more prominent.^5^ Specifically, one large-scale study found evidence for an association between socioeconomic status and diurnal cortisol in children younger than 12 years old, but not in an older sample of 12-16 year-olds.^5^ In line with this proposition, we tested whether a similar pattern could be observed in our data by re-running the same nine regressions in a subsample of children younger than 12 years old. We did not find evidence of a more ubiquitous pattern of associations between measures of the socioecological context and variation in diurnal and pervasive cortisol output in this younger portion of the sample (see Table S7). However, the associations between neighborhood poverty and diurnal cortisol (*β* = 0.349, *p_uncorrected_* = 0.001, *p_corrected_* = 0.0090) and between school-level achievement and pervasive cortisol output (*β* = −0.564, *p_uncorrected_* = 0.001, *p_corrected_* = 0.0090) were characterized by slightly stronger effect sizes, and both remined significant after accounting for false discovery rates.

In a wider age range sample that additionally included high school students, we previously reported that the relationship between family SES and hair cortisol varied by age. ^61^ Therefore, we next tested interactions between each of the nine dimensions of social context and age. There was no evidence of significant interaction effects between age and social context in predicting variation in higher-order factors of HPA function in the current sample (see Table S8). In line with evidence reporting a moderating effect of puberty on the association between cortisol and exposure to disadvantage,^62^ we tested the interaction between the nine dimensions of socioecological context and puberty (see Supplementary Material for a description of the puberty measure adopted). We found no evidence of a significant interaction between puberty and socioecological context in predicting variation in diurnal and pervasive cortisol (see Table S9).

Finally, we conducted a granular analysis of the relationships between each of the seven indices of cortisol output and the nine dimensions of social context (63 pairwise associations). As depicted in Figure 5, while associations were moderate among different indicators of cortisol output and among different dimensions of socioecological context, we did not observe strong or widespread associations between cortisol and socioecological context (Figure 5). Out of the nine dimensions of socioecological context, only three showed significant associations with at least one aspect of cortisol output. First, children attending schools with higher average levels of academic achievement showed steeper declines in cortisol from morning to evening (*r* = −.24, *p_uncorrected_* = .009), such that they produced lower overall levels of cortisol during the day. Second, children in higher-SES neighborhoods had higher cortisol levels at waking (*r* = .23, *p_uncorrected_* = .007) and flatter cortisol awakening responses (*r* = −.28, *p_uncorrected_* = .009). Finally, children whose parents reported more severe interparental conflict had higher in-lab cortisol baseline levels (*r* = 0.13, *p_uncorrected_* = .049).

**Figure 5.**
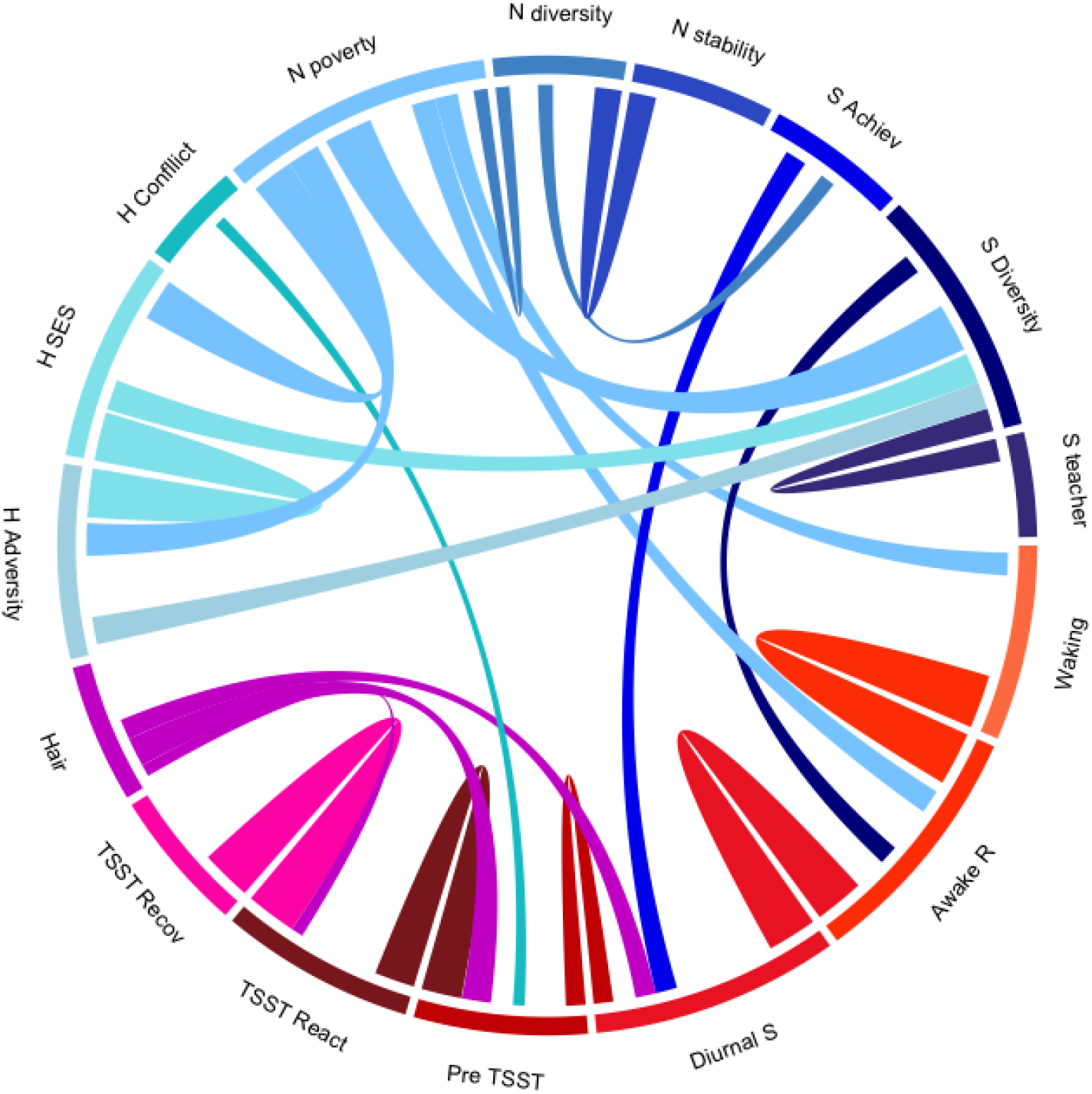
Correlations between multi-modal cortisol output and the nine socioecological context indicators characterizing variation in the home, school and neighborhood environments. The size of each segment corresponds to the proportion of associations that each construct shares with the others relative to the size of associations for every other construct. This technique results in, for example, neighborhood SES, a construct overlapping more substantially with all other measures being represented by a larger segment than, for example, home conflict, a construct weakly related to all others. The size of each river corresponds the size of the correlation between each pair of variables (see supplementary Table S7), all estimates are calculated after accounting for all correlates described in the supplementary information and after including age, sex, age×sex and race as covariates in the model. H = Home, N = Neighborhood, S = School, TSST = Trier Social Stress Test, Diurnal S = Diurnal Slope, Awake R = Awakening Response.

Overall, 4 out of 63 possible pairwise associations between dimensions of social context and indicators of cortisol output were significantly different from zero at a nominal alpha threshold of *p* < .05 (supplementary Table S8). In sensitivity analyses that retained samples that were excluded due to being off-phase with respect to a naturally occurring circadian cortisol rhythm, Supplementary Figure S4, two out of the four nominally significant correlations (parental conflict with in-lab pre-TSST levels and neighborhood SES with awakening response) were no longer observed.

Summarizing across analyses, the only dimension of social context that was reliably associated with cortisol output was concentrated neighborhood poverty, which showed a significant association with a latent factor reflecting diurnal variation in cortisol and which showed nominally significant associations with specific indicators of that factor.

## DISCUSSION

Cortisol is widely studied as a biomarker for the biological embedding of stress. We tested if children exposed to socioeconomic disadvantage in their home, school, and neighborhood contexts showed differences in their cortisol output. There were two main findings. First, measures of cortisol output converged onto two, dissociable dimensions of HPA function. One factor represents pervasive, trait-like accumulations of cortisol; the second represents diurnal change in cortisol. Second, neither dimension of children’s HPA function was strongly or consistently associated with socioecological disadvantage. Rather, the relationship between cortisol output and socioecological disadvantage was specific to a particular HPA function in a particular social context: Children exposed to concentrated neighborhood poverty showed altered diurnal rhythms of cortisol output. No other social context showed a significant association with either major cortisol dimension.

This study is unique in its combination of large sample size, in-depth measurement, and representation of social inequality. Despite being drawn from a circumscribed geographical area, children from the Texas Twin Project vary considerably in their exposure to social disadvantage. Over 30% of families reported having received means-tested public assistance at some point since the children were born),^63^ while the income inequality of the sample rivals levels of inequality seen in countries such as Israel and New Zealand.^56^ We integrated multiple sources of data form multifaceted indicators of social disadvantage over many years. These indices of long-term exposure to social disadvantage were promising candidates as environmental correlates of cortisol output, because HPA axis function is conceptualized as providing a mechanism for long-term adaptation to environmental adversity.^32^ Yet, despite the study being well-suited to detect associations between HPA function and socioeconomic disadvantage, observed associations were generally weak and inconsistent. Most notably, we found no associations between cortisol output and children’s home environments, including conflict between parents, parental socioeconomic status, and cumulative home adversity. These null effects are consistent with some, but by no means all, previous studies.^23,34^

The only association that survived correction for multiple testing was a link between neighborhood disadvantage and diurnal cortisol rhythm. Children living in wealthier neighborhoods showed lower levels of diurnal cortisol, characterized by heightened waking levels, lower awakening response and higher levels maintained throughout the day. This is in line with extant research in adult samples that found that neighborhood poverty was associated with elevated diurnal cortisol levels. ^64^ However, another study found that neighborhood deprivation was associated with lower rates of cortisol recovery after stress exposure in a sample of eighty-five African American children,^65^ a measure of cortisol output that did not emerge as significant in the current work. A specific link between neighborhood concentrated poverty and the biological embedding of stress is consistent with evidence of a specific association between neighborhood deprivation and mortality accounting for many other established socioeconomic risk factors. ^66^ The dysregulation of HPA functioning associated with greater exposure to toxic assault in more deprived neighborhoods^67^ might constitute a potential explanation for the observed link between circadian cortisol dysregulation and neighborhood concentrated poverty. Neighborhood deprivation – nor any other aspect of social disadvantage – was not linked to variation in the pervasive factor of cortisol, which contradicts the perhaps simplistic notion that children exposed to poverty and disadvantage have chronically high levels of cortisol.^68^

In line with evidence of a stronger association between cortisol output and socioeconomic disadvantage in younger samples,^5^ we observed stronger and significant links between pervasive cortisol output and school-level achievement when we conducted our analyses in the younger cohort of children (less than 12 years old). Similarly, the positive association between diurnal cortisol output and neighborhood poverty was characterized by stronger effects. However, even in this younger cohort, we did not find support for a more ubiquitous pattern of association between cortisol output and socioecological disadvantage, as most associations did not reach significance even before correcting for multiple comparisons.

In addition to clarifying the relationship between socioeconomic disadvantage and children’s cortisol output, we also introduced two methodological innovations for the analysis of hormonal data. The first methodological innovation was a multi-level, piecewise latent growth curve modeling approach to analyzing repeated hormonal measurements. We demonstrate how this analytic method can be applied to modeling how people differ in their hormonal change from minute-to-minute, from hour-to-hour, and from day-to-day, while also considering within-person fluctuations. Being able to accurately capture change over time is critical for accurately measuring hormonal function. As described by Shirtcliff et al. (2014, p.44), a fundamental idea for understanding the HPA regulatory system and, more broadly, all biological regulatory systems, is that ‘regulation implies change, fluctuation, and calibration to context’.^68^

The second methodological innovation was a structural equation modeling approach to examine how different aspects of cortisol output converge. As expected, measurements taken on the same timescale (acute responsiveness versus diurnal rhythm) were more strongly related to each other than to measurements taken on different timescales. Measuring cortisol minute-to-minute in the lab, children who showed higher reactivity to an acute stress had lower baseline levels prior to stress exposure and faster recovery following exposure.^38,49^ This well-established pattern is consistent with the proposition that chronically-high cortisol output impairs an individual’s ability to enlist the HPA axis adaptively in times of stress and challenge.^32^ Measuring cortisol hour-to-hour over several days at home, children who had greater awakening responses showed lower waking levels but also maintained higher levels of cortisol throughout the day. Whereas a surge in cortisol secretion in response to an acute stress is potentially adaptive and linked to generally lower levels of pervasive cortisol concentration, an increase in cortisol production soon after waking might be less adaptive and associated with heightened levels of chronic cortisol.

Considering the correlations among cortisol measurements taken at different timescales revealed a dimension of pervasive, trait-like cortisol accumulation, characterized by maintaining higher levels throughout the day, higher levels of cortisol before stress exposure, higher concentrations of cortisol in hair, and blunted reactivity to acute stress in the lab. In contrast, there was less coordination between dynamic aspects of cortisol output: Responsiveness to acute stress and diurnal change were largely unrelated to one another, as has been found by previous research.^69,70^ Overall, the current approach improves upon previous research, which has largely examined pairwise correspondence between individual indices,^49^ and provides new information on the extent to which widely-used research paradigms are tapping convergent versus divergent aspects of HPA function.

Several limitations warrant discussion. The relationship between cortisol output and environmental adversity might be evident only when considering extreme forms of adversity, such as neglect or violence, ^20,21^ which we did not examine. Furthermore, with the exception of interparental conflict, this work focused on aspects of home adversity linked to a family’s wealth and social position, rather than on the emotional stability and support provided by parents. Recent research has started to examine HPA function in relation to positive social interactions and warm, nurturing environments.^71^ This research has shown that stronger attachment, parent-child bonding, and teen-reported positive parenting prospectively predicted higher waking cortisol and steeper diurnal slopes, particularly among Caucasian adolescents.^72^ Animal studies have also suggested that positive environments might be biologically embedded via HPA function.^12,13^ Examining this hypothesis further is an important goal for future research.

Several theories have proposed that associations between cortisol and environmental adversity will be non-linear.^32,33^ Consistent with this idea, we found evidence for a quadratic relationship between school-level achievement and pervasive cortisol factor: Children attending very high and very low-performing schools showed lower levels of trait-like cortisol output, while levels were higher in children attending school characterized by middle levels of achievement. However, the association was not robust after accounting for multiple testing. The difficulty of ascertaining whether the non-linear association with school-achievement was reliably different from zero raises another potential limitation: Associations between environmental disadvantage and cortisol might be more ubiquitous than the current results indicate but might be very small. Even with hundreds to thousands of children, we might be underpowered to detect associations reliably.

In this way, research with hormonal biomarkers might parallel developments in genetic research: Initial enthusiasm about single measurements of a complex biology (candidate genes) gave way to disillusionment about failures to replicate, which ultimately motivated consortia projects that generated sample sizes that were orders-of-magnitude larger than previous studies.^73^ These large sample sizes, coupled with rigorous controls for multiple testing, has resulted in significant breakthroughs in identifying genetic correlates of complex human outcomes. A similar improvement in rigor and sample size might be necessary to advance research using hormonal biomarkers. One possibility would be to form consortia to harmonize the wealth of cortisol data that has already been collected in developmental samples. There will be a particularly strong need for larger sample sizes to powerfully test for individual differences in sensitivity to environmental inputs (*i.e.*, gene × environment interactions).^32^

The difficulty in detecting meaningful associations between socioecological disadvantage and HPA functioning might reflect individual-level heterogeneity in HPA response to environmental exposures. For some HPA output might be up-regulated in response to stress whereas for others the HPA response might be down-regulated. Evidence of heterogeneous changes in hair cortisol secretion in response to a prospective intervention study of cortisol concentration in war-exposed adolescents provides initial support for this possibility.^74^ The randomized 8-week intervention decreased levels of hair cortisol concentration for adolescents characterized by initial *hyper*secretion and medium cortisol secretion, whereas it resulted in increased levels for adolescents starting out with lower levels of cortisol (*hypo*secretion).^74^ More advanced nonlinear and interactive methods may be required to detect these heterogeneous responses of the HPA system in relation to environmental exposure.

## CONCLUSION

We conducted a study of the relationship between cortisol output and social inequality that advances the field by combining depth and comprehensiveness of measurement in a large, population-based sample.^68^ By adopting an in-depth approach to environmental and hormonal measurement, we overcome some of the primary limitations that have characterized previous studies linking exposure to environmental adversity to variation in cortisol output. These previous limitations include using small sample sizes, using single indices of cortisol output or environmental adversity, and adopting variable levels of methodological rigor.^44,68,75,11,48^ Our results showed that selected aspects of neighborhood and school disadvantage were associated with variation in HPA functioning. Contrary to previous reports, we failed to observe a ubiquitous pattern of associations between cortisol output and socioecological disadvantage. Prominent theories position glucocorticoid response as a general biological mechanism for how socioeconomic inequality produces health disparities, but the current results suggest that these theories need to be refined to better account for the specificity of the relationship between cortisol and social context. Much remains to be understood regarding how socioeconomic disadvantage gets under the skin to affect human physiology.^3,76^

## METHOD

### Sample

Participants were members of the Texas Twin Project, a population-based sample of twins and higher order multiple living in Austin, Texas metropolitan area.^77^ Families of twins and other multiples were recruited from public school rosters and invited to take part in the study based at University of Texas at Austin. The University of Texas Institutional Review board granted ethical approval. Participants’ age ranged from 8.06 to 14.75 years (*M* = 10.77, *SD* = 1.83). Sample size varied depending on the cortisol collection modality. A total of 416 samples were available for diurnal variation in cortisol, measured at home over up to five consecutive days per participant. A total of 444 samples were available for cortisol reactivity/recovery to an acute stressor, measured in-lab. A total of 1210 samples were available for hair cortisol concentration, assessing cortisol accumulation over a more extended period of time. Of the total number of unique participants who contributed at-home salivary samples (*n* = 412, 52% females), 400 had also provided in-lab salivary cortisol data (97% coverage), and 382 participants had also contributed hair cortisol data (91.8% sample overlap). The pronouncedly larger sample size for hair samples compared to diurnal and acute stressor samples resulted from the fact that collection of hair samples was introduced into the Texas Twin Project protocol several years before collection of diurnal and acute stressor samples was introduced. The sample was ethnically diverse: 13.4% reported being Hispanic, 64.1% reported being Caucasian, 3.7% reported being African-American, 3.8% reported being Asian-American, 14.4% reported multiple races/ethnicities, and 0.4% reported belonging to other racial/ethnic categories. Supplementary Table S9 to S11 present the sample size available for each measure and the descriptive properties of each variable.

### Measures

#### Salivary Cortisol

##### Diurnal Cortisol Collection At-Home

Participants were instructed to drool passively through a straw into 2 mL plastic vials at three times of day: immediately upon waking, 30 min after waking, and right before bedtime. The median interval between sample1 and sample 3 ranged between 13.65 hours for day 1 and 13.86 hours for day 3 (see supplementary Figure S5a for a visual summary of the diurnal cortisol data collection moments). The vials provided for the at-home salivary collection were color-coded with a different color corresponding to each sampling day, and participants were instructed to place each vial in their home freezer immediately after sampling. Participants were asked to refrain from eating, drinking, or brushing their teeth for the 30 min preceding each sample, and they were provided with diaries where they could record their daily activities and experiences regarding the data collection.

Saliva samples were provided over four consecutive days with a fifth collection day in case of any sampling problem. Seventy participants (16.9%) completed the fifth day of collection in spite of not having experienced any problems. Data coverage for the at-home sampling was excellent, as 378 participants (90.8% of the total sample) completed the first four consecutive days of sampling, and 408 participants (98% of the sample) provided cortisol samples over four days. Participants were instructed to report the date and time (in hours and minutes) of collection by writing on an adhesive label, which they attached to each vial after sampling. In order to assess time reporting accuracy, each sampling vial had to be removed from a bottle equipped with a date and time-tracking cap (MEMs Track Cap; Aardex, Denver, CO). The slight deviation between the reported sampling time and the time stamp derived from the MEMs cap intuitively indicated sampling duration, the median deviation time ranged between 3 and 4 minutes for the five sampling days. Saliva samples and study materials were returned to the lab using a pre-paid envelope provided to every family as part of the at-home collection kit, the day after saliva collection was completed. Samples were frozen at −40 degrees in the laboratory prior to being shipped on dry ice to Dr. Clemens Kirschbaum’s laboratory in Dresden, Germany for assay using liquid chromatography tandem mass spectrometry (LC-MS/MS).

##### Reactivity/Recovery Cortisol Collection In-Laboratory

To examine the hormonal signature of responses to an acute stress, participants were asked to participate in the Trier Social Stress Test (TSST; ^38^) during their visit to the research laboratory. After approximately 30 min from their arrival to research lab, participants were taken to a different space and instructed to prepare a short story to present to an unfamiliar audience comprising two “judges.” After the story preparation (3 min) and presentation time had elapsed (5 min), participants were instructed to perform mental arithmetic in front of the judges (5 min). Although twin pairs and triplets came to the research laboratory together, each child completed the TSST separately. Of the 444 individuals that participated in the TSST, 85 discontinued the task before completion but provided post-Trier saliva samples when willing. Four cortisol samples were collected to examine participants’ response to an acute, standardized stressor: The first sample was collected upon arrival to the laboratory, at least 30 minutes before the TSST; The second sample was collected 20 minutes after the start of the TSST; The third sample was collected 20 minutes after the completion of Sample 2; The fourth sample was collected 20 minutes after the completion of Sample 3 (see Figure S5b for a visual summary of the in-lab cortisol data collection moments). Participants were instructed to refrain from eating one hour prior their visit to the research laboratory. Participants contributed their samples by drooling passively through a straw into a 2ml plastic vial, and the research assistant helping with the research visit recorder the exact time at which each sample was collected. All samples were frozen at the same time, within maximum two and a half hours from the collection of the first sample, at −40 degrees prior to being shipped on dry ice to Dr. Clemens Kirschbaum’s laboratory for assay using liquid chromatography tandem mass spectrometry (LC-MS/MS).

#### Hair Cortisol

During the research visit, research assistants collected hair samples from 1210 participants (92.5% of all in-lab visits as of spring 2018). Samples were not available for the remaining observations due to either their hair being too short, or participants having declined to provide a sample. Although hair hormones are robust to a number of possible confounds, including hair products and wash frequency ^78^, participants were instructed to refrain from using leave-in hair products, such as hair gel, on the day of the lab visit. Hair samples of approximately 3 mm in diameter and at least 3 cm in length were obtained from the posterior vertex position (i.e., the center of the back of the head). The 3-cm hair segment closest to the scalp was analyzed as a marker for average cortisol secretion over the most recent 3-month period. Samples were cut as close to the scalp as possible from the center of the back of the head, stored in a dry location, and shipped to the Technical University of Dresden for steroid extraction and measurement (technical details on the extraction procedure are provided elsewhere ^79^). Internal consistency estimates for cortisol analyzed using liquid chromatography tandem mass spectrometry (LC-MS/MS) have been reported above .96.^80^ In a sub-sample of participants from the Texas Twin Project, reliability for cortisol samples analyzed in duplicate was estimated at .89, and concordance between monozygotic twins was found to be high.^81^ The lower limit of sensitivity for hair cortisol was 0.1 pg/ml.^79^

Quality control processes and exclusion criteria for all cortisol modalities are reported in detail in the Supplementary Methods.

#### Socioecological contexts

Multiple indices of adversity and socioecological deprivation were calculated for home, school, and neighborhood contexts (see Engelhardt et al., 2018 for a detailed description). Three indices were created for the <u>*home environment*</u>: (a) Parent socio-economic status, obtained from a standardized composite of parent reported income (log transformed) and educational attainment; (b) Cumulative adversity, which was created by averaging eight variables that measured the presence or absence of financial difficulty during the twins’ lifetime, as well as major life changes in the six years preceding the twins’ study participation (self-reported food security, public assistance, changes in home address, income, parental education and occupation, history of financial problems and father absence); (c) Parent conflict, measured with the Porter & O’Leary’s scale (1980) which assessed children’s exposure to at-home conflict related to finances and discipline.^82^

Three additional indices were created for the <u>*neighborhood environment*</u>. These were constructed using multiple indices available through the American Community Survey, an annual survey administered by the U.S. Census Bureau to gather information on resident demographics, employment, and housing characteristics (United States Census Bureau). Data for the variable of interest for years 2011-2017 for each of the 239 census tracts in which the current sample’s participants resided. Tract-specific estimates for each of the 12 variables of interest were averaged across available years to generate cross-year indicators of neighborhood quality for every tract. These average estimates were submitted to a series of Principal Component Analyses (PCAs) to generate weights. Consequently, the year-specific data weighted by the corresponding unstandardized loadings derived from the PCA, and weighted composite scores were constructed for each of the three indicators: (a) Neighborhood concentrated poverty (created combining educational attainment, single motherhood, management positions, impoverishment, and unemployment); (b) Residential stability (created combing information about: housing owned, relocation in the past year, maintain the same residence for a decade, and number of children and adolescents); and (c) Neighborhood diversity (created from a weighted composite of racial/ethnic minority status and immigration).

Finally, three indices created for the <u>*school environment*</u>. These were derived from yearly state-mandated reports by the Texas Education Agency. Similarly to the neighborhood data, the school composited were derived combining estimates for each variable of interest across available years (2011-2017), submitting these cross-year indicators to PCAs, and creating weighted composite scores indexing three characteristics: (a) school-level achievement (attendance, as well as proficiency on a statewide test of math and reading); (b) student demographics (students’ racial/ethnic minority status, English language learner status, low SES by virtue of eligibility for free/reduced lunch, and mobility); and (c) teacher characteristics (years of teaching experience, salary, and student-to-teacher ratio). A detailed description of the procedure is provided in Engelhardt et al.^53^

### Analytic Strategies

#### Data transformations and estimation

Cortisol levels were log transformed prior analyses to correct for positive skew, descriptive statistics for the log transformed scores are reported in supplementary Table S8 and distributions shown in Figure S6a-c. Distributions for the socio-ecological indicators are shown in Supplementary Figure S6d and descriptive statistics in Table S9. All models were fit with full information maximum likelihood estimation, which accommodates uneven patterns of missingness under the assumption that, conditional on the observed data that are included in the model, the pattern of missingness is unrelated to the missing values. As individual participants in this sample were nested within families, non-independence of observations was accounted for by applying a sandwich estimator, specified in MPlus syntax as TYPE=COMPLEX, in all analyses reported in the current work. Additional details on the exclusion criteria, controls and covariates are provided in the Supplementary Information.

#### Piecewise Latent Growth Curve Models: A Novel Approach for Characterizing Children’s Cortisol Output over Time

The statistical approaches that have been adopted to modeling individual variation in diurnal cortisol take into account its naturally occurring daily fluctuation. One of such approaches entails calculating the area under the curve (AUC; Fekedulegn et al., 2007; Pruessner, Kirschbaum, Meinlschmid, & Hellhammer, 2003). A further approach, which considers the normative pattern of diurnal cortisol fluctuation, has been applying latent growth models to model of variation in cortisol patterns.^49,59^ In the current work, we applied multi-level latent growth models to better capture the change in salivary cortisol at the intra-and inter-individual levels. Mplus version 8.0 was used to conduct all the multi-level growth modelling.^83^

To model individual differences in the <u>diurnal cortisol</u> trajectories while accounting for inter-individual variability, we initially applied a three-level latent growth model, where Level 1 captured the within person cortisol trajectory, Level 2 denoted day-to-day variation across the five sampling days, and Level 3 described the between person variation after having accounted for the within-person and day-to-day variation. Testing this three-level approach, we observed that the day-to-day variation explained a very small proportion of variance, consequently we opted for a two-level latent growth model approach to best, and most parsimoniously, capture diurnal cortisol variation. Within this two-level growth model framework, Level 1 represented within-person variation in the diurnal cortisol trajectory and Level 2 denoted between-person variation actor controlling for the effect of intra-individual variability, in addition the day-to-day effect was accounted for by including days as dummy coded covariates at Level 1.

At Level 1, we specified a piecewise latent growth model approach including one latent factor for the intercept, reflecting variation in initial cortisol levels; a latent slope, representing variation in cortisol awakening response, and a second latent slope, representing variation in the naturally occurring decline in cortisol level throughout the day. Each latent factor constituted a random effect and was consequently allowed to vary at Level 2 (between participants).

The Level 1 model representing within-person cortisol trajectories in diurnal cortisol and can be expressed as:

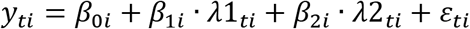

where *y*_*ti*_ represents cortisol values at sampling time *t* for individual *i*; *β*_0*i*_ represents initial cortisol levels for individual *i*; *β*_1*i*_ and *β*_2*i*_ represent the magnitude of rise and decline in cortisol prior to and after the turning point, respectively; *λ*1_*ti*_ and *λ*2_*ti*_ represent the time-specific basis coefficients defining the rise and decline of cortisol, respectively. We applied a data-driven approach to determine the basis coefficients for modelling the turning point for both diurnal and reactive cortisol trajectories, which is described in the following section. Finally, *ε*_*ti*_ represents a sample-specific residual variance for an individual that is modelled at Level 2. The Level 2 model for between-person effects can be expressed as the following equations:

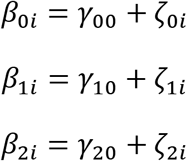

where *γ*_00_ is the average initial cortisol level across individuals, *γ*_10_ is the mean rise across individuals prior to the turning point, *γ*_20_ is the mean decline across individuals after the turning point, and the *ζ* terms represent person-specific deviations from the mean.

This two-level growth approach was used to model both diurnal cortisol output and in <u>reactivity/recovery in cortisol output in response to the TSST</u>. Within the diurnal model, *β*_0*i*_ is a random intercept reflecting variation in waking cortisol levels, *β*_1*i*_ is a latent slope describing the cortisol awakening response, and *β*_2*i*_ is a second latent slope representing variation in the diurnal slope from morning to evening. Within the TSST model, *β*_0*i*_ is a random intercept reflecting variation in initial cortisol levels (measured upon arrival to the research laboratory), *β*_1*i*_ is a latent slope describing the in the initial rise in cortisol following the TSST (reactivity to an acute stress), and *β*_2*i*_ is a second latent slope representing variation in the subsequent decline in cortisol following the TSST test (recovery after an acute stressor).

We adopted a data driven approach to determine the turning point to apply to the two-level piecewise latent growth models. In order to estimate the location of the turning point, we fit a series of models in which the slopes coefficients for cortisol awakening response (and the equivalent cortisol reactivity following the TSST) and cortisol decline varied as function of individuals’ sampling times relative to a range of possible turning points.

With respect to at-home saliva samples, the diurnal cortisol awakening response has been shown to peak between 30 and 45 minutes after wake, followed by a decline over the course of the day. In line with this normative diurnal pattern, participants were instructed to provide the second cortisol sample 30 minutes after waking; however, there was individual variation around the time of the all sampling moments. We leveraged the variation in sampling times to determine the true turning point at which salivary cortisol peaked and began its decline following the awakening response. To do this, we fit a series of multilevel latent growth models in which the basis coefficients of the latent slopes varied as function of individuals’ sampling times relative to possible turning points. We tested turning points in 1-min increments from 25 to 40 minutes from waking time and compared the model fit for each growth curve to establish the optimal data-driven turning point. The best fitting model, based on the Log Likelihood and AIC values, was one in which a turning point of 32 min after waking time (see Supplementary Figure S7a). This parametrization, with a turning point at 32 minutes after waking, was therefore applied in all subsequent analyses. Additional information on this data-driven approach to estimate turning points is provided in the Supplementary Methods.

A similar approach was adopted to test the optimal turning point for the in-lab reactivity/recovery in cortisol trajectory, with the only difference that the time interval prior the start of the TSST was fixed to 0, so as to identify a random intercept representing basal, pre-stressor, cortisol levels. The optimal turning point was found to be 25 minutes from the start of the TSST (see Supplementary Figure S7b). This parameterization, with a turning point at 25 minutes after start of the TSST, was adopted in all subsequent analyses.

## Author contributions

MM, KPH, and EMT designed the study; MM, LEE, KPH, and EMT contributed new reagents/analytic tools; MM, LEE, and EMT analyzed data; MM, LEE, LR, AS, ADG, DAB, JWM, SMF, MWP, KPH, and EMT performed the research; and MM, KPH, and EMT wrote the paper.

The authors declare no conflict of interest.

## Acknowledgements

We gratefully acknowledge all participants members of the Texas Twin Project. KPH and EMT are Faculty Research Associates of the Population Research Center at the University of Texas at Austin, which is supported by a grant, 5-R24-HD042849, from the Eunice Kennedy Shriver National Institute of Child Health and Human Development (NICHD). KPH and EMT are also supported by Jacobs Foundation Research Fellowships. This research was supported by NIH grant R01HD083613. MM is partly supported by the David Wechsler Early Career Award for Innovative Work in Cognition and by NIH grant R01HD083613 awarded to ETD and KPH.

## Code Availability Statement

Results, Full Information Maximum Likelihood (FIML) summary data, analytic scripts, and generated outputs will be uploaded and instantly available for all researchers to use.

## Data Availability Statement

While results, FIML summary data, analytic scripts, and generated outputs will be uploaded and instantly available for all researchers to use, our policy regarding the access of raw data files is separate. The data file related to this project contains particularly sensitive information, including each child’s geocoded neighborhood and school information and sensitive endocrinological data. To this end, researchers will be able to obtain the data file through managed access. Requests for managed access should be sent to Dr. Elliot Tucker-Drob and Dr. Paige Harden, joint principal investigators of the Texas Twin Project.

