## Supplementary Information for "Weak and uneven associations of home, neighborhood and school environments with stress hormone output across multiple time scales"

### Supplementary Methods

- Quality control, exclusions and covariates applied to cortisol data
- Data-driven approach to determining salivary cortisol turning points (TPs) for piecewise growth models.

### Supplementary Tables

- **Table S1.** Two-level latent growth model capturing variation in diurnal cortisol.
- **Table S2.** Two-level latent growth model capturing variation in reactive cortisol.
- **Table S3.** Two-level latent growth model of parallel processes, capturing variation in diurnal, reactive and hair cortisol.
- **Table S4.** Two-level latent growth model of parallel processes, capturing variation in diurnal, reactive and hair cortisol including second-order cortisol factors
- **Table S5.** Associations between the nine sociecological context indicators and the two second order latent factors of 'pervasive' and 'diurnal' cortisol output.
- **Table S6.** Associations between the quadratic terms for the nine sociecological context indicators and the two second order latent factors of 'pervasive' and 'diurnal' cortisol output.
- **Table S7.** Associations between e sociecological context indicators and the two second order latent factors of 'pervasive' and 'diurnal' cortisol output in the younger portion of our sample (< 12 years).
- **Table S8.** Associations between e sociecological context indicators and the two second order latent factors of 'pervasive' and 'diurnal' cortisol output by age.
- **Table S9.** Associations between e sociecological context indicators and the two second order latent factors of 'pervasive' and 'diurnal' cortisol output by puberty.
- **Table S10.** Correlations between all cortisol indicators and the nine socioecological context indicators.
- **Table S11.** Final sample size after basic exclusions for diurnal, reactive and hair cortisol.
- **Table S12.** Descriptive statistics for log transformed diurnal, reactive and hair cortisol samples.
- **Table S13.** Descriptive statistics for socio-ecological context measures.

### Supplementary Figures

- **Figure S1.** Correlations between the nine socioecological context indicators.
- **Figure S2.** Intra and inter-individual variability in diurnal cortisol levels over the five sampling days.
- **Figure S3.** Regression line for the association between pervasive cortisol output and the quadratic term for school-level achievement
- **Figure S4.** Associations between cortisol latent factors and SES context measures in the final sample (panel a) and in the full sample, including participants showing off-phase CORT rhythm (panel b).
- **Figure S5.** Timeline of data collection procedure for (a) in-home diurnal cortisol and (b) in-lab reactive cortisol.
- **Figure S6.** Distributions of (**a**) log-transformed at-home diurnal cortisol samples, (**b**) log transformed in-lab reactive cortisol samples, (**c**) log transformed hair cortisol values, and **(d)** socio-ecological context measures.

- **Figure S7**. Model fit indices comparison for alternative latent growth models considering a range of turning points for (a) diurnal cortisol and (b) reactive cortisol trajectories.
- **Figure S8.** Longitudinal coverage and level of log-transformed at-home cortisol samples after all exclusions over the 5-day collection period (a), and reintroducing the cases excluded for being off-phase (b).

**Quality control, exclusions and covariates applied to cortisol data**

***Diurnal cortisol output.*** MEMs cap times were unavailable for 13.30% of returned samples, primarily due to product failure or because participants had opened the MEMs container for samples 1 and 2 simultaneously. For those with MEMs data available, the mean deviation of MEMs recorded time from that reported by participants was 6.19 minutes across all samples and days. Consequently, participants and parents reported sampling times were used for the current quality control procedures and following analyses. Based on recommended criteria for analyzing cortisol data,[1] participants were excluded from all analyses if they were taking steroid-based medication, if they were on hormone-based contraceptives, and if they reported hormone-related dysfunctions. A well-documented normative pattern characterizes the daily production of cortisol with cortisol levels increasing in the second half of the sleep cycle, peaking between 30 to 45 minutes after waking, and then steadily declining throughout the day [2,3]. In order to identify individual variation in this normative pattern of activity, the first cortisol sample was collected every day upon waking and the second sample 30 minutes after the first one. In line with this normative pattern of assessment, cases were excluded from analyses when the time interval between sample 1 and 2 exceeded 60 minutes (daily $N$ ranged from 5 to 14 across the 5 days of data collection). Samples were also excluded when the interval between wake and sample 1 exceeded 20 minutes, as considered not indicative of waking levels, but of the cortisol awakening response ($N$ ranging from 0 samples for day 3 and 6 for day 1). At-home samples were also excluded when cortisol concentrations for the wake+30 sample were lower than concentrations for the evening sample on the same day or when concentrations for the evening sample were higher than the next day's waking concentrations ($N$ excluded day1 = 34, $N$ excluded day 2 = 31, $N$ excluded day 3 = 28, $N$ excluded day 4 = 38, $N$ excluded day 5 = 16). These conditions suggest either a failure to provide saliva at the correct sampling times, or an abnormal wake-sleep schedule. Figure S8a provides an outline of the data coverage for diurnal cortisol over the five sampling days and the levels of CORT concentration for each participant at each sampling moment. Figure S8b provides an outline of the data coverage and diurnal CORT output for the full sample prior to excluding the samples showing off-phase values. In addition to these quality control and exclusion procedures, five participants had contributed diurnal data over two collections, these data were included in the analyses controlling for nestedness within person at each collection wave as well as within family. The remaining sample sizes after all basic exclusions are reported in Supplementary Table S9 and S10. Cortisol values for the remaining samples were log-transformed to correct for positive skew. As recommended by published guidelines,[1] we controlled for several potential confounders by including these variables as covariates in the reported analyses and accounting for their effect by means of linear regression. In dealing with at-home diurnal CORT samples, the following covariates were considered: (1) assay batch, (2) variation in waking time, (3) Reported dairy intake, (4) food intake between sample 1 and sample 2, (5) non-steroid medications taken on the day of sampling.

***Reactivity/Recovery cortisol output.***

The same basic exclusion criteria that we applied for the diurnal CORT samples were applied to the in-lab reactivity/recovery CORT data, and a total of 17 participants (3.8%) were excluded from analyses due to taking steroid-based medication, if they were on hormone-based contraceptives, and if they reported hormone-related dysfunctions. The remaining sample sizes are reported in Supplementary Table S10. Cortisol values for the remaining samples were log-transformed to correct for positive skew. As for the diurnal salivary samples, we accounted for several potential confounders by including a series of covariates in the reported analyses. When modelling in-lab cortisol data the following covariates were included in the model: (1) assay batch, (2) non-steroid based medications taken up to two weeks prior the in-lab appointment, (3) non-steroid based medication taken on the day of the in-lab appointment, (4) time between waking and sampling times.

***Hair cortisol***

Hair samples were excluded when participants had a hormone-disrupting disorder (5 samples), or when participants had taken steroid-based medication regularly in the past six months (9 samples). Cortisol values for the remaining samples (Table S8) were log-transformed to correct for positive skew and residualized for assay batch, which was significantly related to cortisol variability ($F$ (3, 145) = 20.21, $p$ < .0001). We tested for associations between hair cortisol and other potential confounds, including hair care routines, whether their hair was altered chemically and whether they had hair extensions were also considered. Of these potential confounders, significant negative associations with hair cortisol were found when participants had dyed or highlighted hair (7%; $t$ (142) = -2.30, $p$ < .05) and when they reported blow drying their hair regularly (25%; $t$ (142) = -2.10, $p$ < .05). A significant positive association occurred between hair cortisol and use of hair extensions (<1%; $t$ (503) = 2.84, $p$ < .01). The effect of these confounders was statistically accounted for my means of multiple regression and the residual variance standardized.  Several participants had contributed hair samples twice, as for diurnal cortisol samples, we included the data in our analyses by controlling for nestedness within person at each collection wave as well as within family.

**Data-driven approach to determining salivary cortisol turning points (TPs) for piecewise growth models.**

We adopted a data driven approach to determine the turning point to apply to the two-level piecewise latent growth models. Such a model estimates a slope prior to a specified turning point and an independent slope reflecting the trajectory following the turning point. In order to converge upon the true turning point observed in the data, we fit a series of models in which the slope basis coefficients for cortisol awakening response (and the equivalent cortisol reactivity following the TSST) and CORT decline varied as function of individuals' sampling times relative to a range of possible turning points.

With respect to at-home saliva samples, the diurnal CORT awakening response has been shown to peak between 30 and 45 minutes after wake, followed by a decline over the course of the day. In line with this normative diurnal pattern, we asked our participant to provide the second cortisol sample 30 minutes after waking; however, there was individual variation around the time of the all sampling moments. We leveraged the variation in sampling times to determine the true turning point at which salivary CORT peaked and began its decline following the awakening response. To do this, we fit a series of multilevel latent growth models in which the basis coefficients of the latent slopes varied as function of individuals' sampling times relative to possible turning points. We tested turning points, $TP$, in 1-min increments from 25 to 40 minutes from waking time and compared the model fit for each growth curve to establish the optimal data-driven turning point. Time was scaled in minutes relative to the tested turning point and then rescaled in hours. (i.e. $time_s/60$) at the modelling stage. Within the multilevel modelling framework repeated samples ($s$) are organized in long format and the basis coefficients for the determination of the turning point in the latent slopes were modelled as follows:

$$\text{If } time_s \leq TP, \text{ then } \lambda 1_s = \frac{time_s}{60}$$
$$\text{If } time_s > TP, \text{ then } \lambda 1_s = \frac{TP}{60}$$
$$\text{If } time_s \leq TP, \text{ then } \lambda 2_s = 0$$
$$\text{If } time_s > TP, \text{ then } \lambda 2_s = \frac{time_s - TP}{60}$$

The best fitting model, based on -2 Log Likelihood and AIC values, was one in which a turning point of 32 min after waking time (see Supplementary Figure S7a).

A similar approach was adopted to test the optimal turning point for the in-lab reactivity/recovery in CORT trajectory, with the only difference that the time interval prior the start of the TSST was fixed to 0, which is reflected as follows:

$$\text{If } time_s \leq 0, \text{ then } \lambda 1_s = 0$$

CORT levels are expected to peak approximately 20 min after the beginning of the TSST, followed by a decline in the 40 min following the peak, as CORT concentration tends to return to the baseline levels.[4] Consequently, we fit a series of multilevel latent growth models in which the basis coefficients of the latent slopes representing cortisol reactivity and decline varied as function of individuals' sampling times relative to turning points ranging from 10 to 35 minutes following the start of the TSST. The optimal turning point was found to be 25 minutes from the start of the TSST (see Supplementary Figure S7b).

***Multi-level latent growth models of parallel processes.***

After modelling variation in diurnal and reactive cortisol independently, we jointly modelled the diurnal and reactive cortisol trajectories using a parallel processes approach. Latent growth modelling of parallel processes is specifically suited to answer questions of how the trajectories of two systems are related to one another.[5] Consequently, we modelled diurnal and reactivity/recovery trajectories in parallel in order to examine the correspondence between these CORT formats. Within this multivariate multi-level approach, at Level 1 we specified within-individual trajectories in diurnal and reactive CORT in the same way described in the previous section. At Level 2 we examined the covariance between the two latent intercepts (one for diurnal and one for reactive CORT) and four slopes (two for diurnal and two for reactive CORT).

Within this two-level growth modelling framework, it is possible to add correlates at either level of analyses (intra- or inter-individual). We added hair CORT as a correlate at Level 2 in order to assess the correspondence between variation in hair CORT concentration and diurnal and reactive CORT output having controlled for intra-individual variability in their trajectories at Level 1. In the same way, age, sex, race, and the socioecological context indicators were included as Level 2 correlates.

**Table S1.** Two-level latent growth model capturing variation in diurnal cortisol

| | Measures | Estimate | S.E. | p |
|---|---|---|---|---|
| ***Level 1: Within-Person Variation*** | | | | |
| | Waking Levels →DAY2 | 0.040 | 0.031 | 0.197 |
| | Waking Levels →DAY3 | 0.033 | 0.030 | 0.261 |
| | Waking Levels →DAY4 | 0.066 | 0.037 | 0.075 |
| | Waking Levels →DAY5 | 0.102 | 0.049 | 0.037 |
| Residual Variance | Waking Levels | 0.345 | 0.024 | 0.000 |
| ***Level 2: Between-Person Variation*** | | | | |
| Covariances Growth Parameters | Waking Levels (intercept) ⇔ Awakening response (slope 1) | -0.213 | 0.111 | 0.054 |
| | Waking Levels (intercept) ⇔ Diurnal Slope (slope 2) | -0.006 | 0.004 | 0.126 |
| | Awakening Response (slope1) ⇔ Diurnal Slope (slope 2) | 0.016 | 0.007 | 0.015 |
| Means | Waking Levels | 1.920 | 0.046 | 0.000 |
| | Awakening Response | 0.724 | 0.074 | 0.000 |
| | Diurnal Slope | -0.213 | 0.004 | 0.000 |
| Variances | Waking Levels | 0.244 | 0.068 | 0.000 |
| | Awakening Response | 0.527 | 0.202 | 0.009 |
| | Diurnal Slope | 0.002 | 0.000 | 0.000 |
| **Model Fit Indices:** -2Log Likelihood = -4329.173, AIC = 8686.347 | | | | |

***Note.*** All estimates are unstandardized; → = regression path; ⇔ correlation path.

**Table S2.** Two-level latent growth model capturing variation in reactive cortisol

| | | | | |
|---|---|---|---|---|
| | ***Level 1: Within-Person Variation*** | | | |
| | **Measures** | **Estimate** | **S.E.** | **p** |
| Residual Variance | Pre-TSST Levels | 0.102 | 0.011 | 0.000 |
| | ***Level 2: Between-Person Variation*** | | | |
| Covariances Growth Parameters | Pre-TSST Levels (intercept) ⇔ TSST Reactivity (slope 1) | -0.692 | 0.120 | 0.000 |
| | Pre-TSST Levels (intercept) ⇔ TSST Recovery (slope 2) | -0.027 | 0.045 | 0.548 |
| | TSST Reactivity (slope1)⇔ TSST Recovery (slope2) | -0.755 | 0.129 | 0.000 |
| Means | Pre-TSST Levels | 0.764 | 0.051 | 0.000 |
| | TSST Reactivity | 1.031 | 0.134 | 0.000 |
| | TSST Recovery | -0.670 | 0.053 | 0.000 |
| Variances | Pre-TSST Levels | 0.667 | 0.127 | 0.000 |
| | TSST Reactivity | 3.811 | 0.394 | 0.000 |
| | TSST Recovery | 0.455 | 0.081 | 0.000 |
| | **Model Fit Indices:** -2Log Likelihood = -1399.641, AIC = 2819.282 | | | |

***Note.*** All estimates are unstandardized; ⇔ correlation path.

**Table S3.** Two-level latent growth model of parallel processes, simultaneously capturing variation in diurnal, reactive and hair cortisol

| | | | | | |
|---|---|---|---|---|---|
| | ***Level 1: Within-Person Variation*** | | | | |
| | **Measures** | **Estimate** | **S.E.** | **p** | |
| | Waking Levels →DAY2 | 0.040 | 0.031 | 0.201 | |
| | Waking Levels →DAY3 | 0.033 | 0.030 | 0.270 | |
| | Waking Levels →DAY4 | 0.065 | 0.037 | 0.079 | |
| | Waking Levels →DAY5 | 0.105 | 0.049 | 0.0344 | |
| Residual Variance | Pre-TSST Levels | 0.102 | 0.011 | 0.000 | |
| Residual Variance | Waking Levels | 0.346 | 0.024 | 0.000 | |
| | ***Level 2: Between-Person Variation*** | | | | $p_{corrected}$ |
| Covariances Growth | Waking Levels ⇔ Awakening response | -0.216 | 0.110 | 0.051 | |
| Parameters | | | | | 0.134 |
| | Waking Levels ⇔ Diurnal Slope | -0.006 | 0.004 | 0.127 | 0.234 |
| | Waking Levels ⇔ Pre-TSST Levels | 0.109 | 0.063 | 0.082 | 0.191 |
| | Waking Levels ⇔ TSST Reactivity | -0.081 | 0.089 | 0.358 | 0.501 |
| | Waking Levels ⇔ TSST Recovery | -0.018 | 0.022 | 0.420 | 0.551 |
| | Waking Levels ⇔ hair concentration | 0.050 | 0.077 | 0.513 | 0.601 |
| | Awakening Response ⇔ Diurnal Slope | 0.016 | 0.007 | 0.017 | 0.060 |
| | Awakening Response ⇔ Pre-TSST Levels | 0.026 | 0.053 | 0.622 | 0.622 |
| | Awakening Response ⇔ TSST Reactivity | 0.084 | 0.164 | 0.608 | 0.622 |
| | Awakening Response ⇔ TSST Recovery | -0.025 | 0.049 | 0.608 | 0.622 |
| | Awakening Response ⇔ hair concentration | 0.100 | 0.069 | 0.145 | 0.234 |
| | Diurnal Slope ⇔ Pre-TSST Levels | 0.008 | 0.003 | 0.004 | 0.021 |
| | Diurnal Slope ⇔ TSST Reactivity | -0.012 | 0.008 | 0.131 | 0.234 |
| | Diurnal Slope ⇔ TSST Recovery | 0.003 | 0.003 | 0.323 | 0.485 |
| | Diurnal Slope ⇔ hair concentration | 0.014 | 0.004 | 0.000 | 0.000 |
| | Pre-TSST Levels ⇔ TSST Reactivity | -0.688 | 0.119 | 0.000 | 0.000 |
| | Pre-TSST Levels ⇔ TSST Recovery | -0.029 | 0.044 | 0.515 | 0.601 |
| | Pre-TSST Levels ⇔ hair concentration | 0.300 | 0.129 | 0.020 | 0.060 |
| | TSST Reactivity ⇔ TSST Recovery | -0.751 | 0.129 | 0.000 | 0.000 |

| | | | | | |
|---|---|---|---|---|---|
| | TSST Reactivity ⇔ hair concentration | -0.363 | 0.137 | 0.008 | 0.034 |
| | TSST Recovery ⇔ hair concentration | 0.076 | 0.052 | 0.140 | 0.234 |
| Means | Waking Levels | 1.921 | 0.046 | 0.000 | |
| | Awakening Response | 0.720 | 0.074 | 0.000 | |
| | Diurnal Slope | -0.213 | 0.004 | 0.000 | |
| | Pre-TSST Levels | 0.763 | 0.051 | 0.000 | |
| | TSST Reactivity | 1.035 | 0.134 | 0.000 | |
| | TSST Recovery | -0.668 | 0.053 | 0.000 | |
| | Hair concentration | -0.014 | 0.071 | 0.841 | |
| Variances | Waking Levels | 0.243 | 0.068 | 0.000 | |
| | Awakening Response | 0.528 | 0.201 | 0.009 | |
| | Diurnal Slope | 0.002 | 0.000 | 0.000 | |
| | Pre-TSST Levels | 0.664 | 0.125 | 0.000 | |
| | TSST Reactivity | 3.805 | 0.395 | 0.009 | |
| | TSST Recovery | 0.453 | 0.081 | 0.000 | |
| | Hair concentration | 1.376 | 0.169 | 0.000 | |
| **Model Fit Indices:** -2Log Likelihood = -6289.920, AIC = 12661.841 | | | | | |

***Note.*** All estimates are unstandardized; → = regression path; ⇔ correlation path, $p_{corrected}$ = p value estimated after correcting for multiple testing using the Bejamini-Hochberg FDR method.

**Table S4.** Two-level latent growth model of parallel processes, capturing variation in diurnal, reactive and hair cortisol including second-order cortisol factors: Factor loadings

| Factor | CORT measures | Estimate | SE | p |
|---|---|---|---|---|
| Diurnal factor | Waking Levels | -0.998 | 0.001 | 0.000 |
| | Awakening Response | 0.559 | 0.174 | 0.001 |
| | Diurnal Slope | 0.516 | 0.193 | 0.007 |
| Pervasive factor | Hair concentration | 0.474 | 0.097 | 0.000 |
| | Diurnal Slope | 0.699 | 0.210 | 0.001 |
| | Pre-TSST Levels | 0.663 | 0.104 | 0.000 |
| | TSST Reactivity | -0.236 | 0.073 | 0.001 |
| | TSST Reactivity ⬄ TSST Recovery | -0.793 | 0.112 | 0.000 |
| | Pre-TSST Levels ⬄ TSST Reactivity | -0.475 | 0.123 | 0.000 |
| | Diurnal factor ⬄ Pervasive factor | -0.394 | 0.179 | 0.028 |
| | Model Fit Indices: -2Log Likelihood = -6289.920, AIC = 12661.841 | | | |

**Note:** the estimates are standardized, ⬄ = correlation path.

**Table S5.** Associations between the nine socioecological context indicators and the two second order latent factors of 'pervasive' and 'diurnal' CORT output.

| Measures | *β* | *p* |
|---|---|---|
| Diurnal factor ⇔ Pervasive factor | -0.453 | 0.066 |
| Diurnal factor ⇔ Home Conflict | -0.089 | 0.135 |
| Pervasive factor ⇔ Home Conflict | 0.133 | 0.074 |
| Diurnal factor ⇔ Home SES | 0.092 | 0.195 |
| Pervasive factor ⇔ Home SES | 0.126 | 0.202 |
| Diurnal factor ⇔ Home Cumulative Adversity | 0.088 | 0.253 |
| Pervasive factor ⇔ Home Cumulative Adversity | 0.066 | 0.396 |
| Diurnal factor ⇔ School-level Achievement | -0.062 | 0.081 |
| Pervasive factor ⇔ School-level Achievement | -0.045 | 0.707 |
| Diurnal factor ⇔ School Diversity | -0.065 | 0.353 |
| Pervasive factor ⇔ School Diversity | 0.071 | 0.407 |
| Diurnal factor ⇔ School Teacher Quality | 0.031 | 0.566 |
| Pervasive factor ⇔ School Teacher Quality | 0.098 | 0.237 |
| **Diurnal factor ⇔ Neighborhood Poverty** | **-0.190** | **0.002** |
| Pervasive factor ⇔ Neighborhood Poverty | -0.012 | 0.885 |
| Diurnal factor ⇔ Neighborhood Diversity | 0.027 | 0.57 |
| Pervasive factor ⇔ Neighborhood Diversity | 0.021 | 0.79 |
| Diurnal factor ⇔ Neighborhood Instability | 0.072 | 0.323 |
| Pervasive factor ⇔ Neighborhood Instability | -0.12 | 0.223 |

**Table S6.** Associations between linear and **quadratic terms** for the nine socioecological context indicators and the two second order latent factors of 'pervasive' and 'diurnal' cortisol output.

| Measures | *estimate* | *p* | *Estimate²* | *p* |
|---|---|---|---|---|
| Diurnal factor⇔ Pervasive factor | 0.414 | 0.029 | - | - |
| Age→ Diurnal | 0.031 | 0.556 | - | - |
| Age→ Pervasive | 0.041 | 0.503 | - | - |
| Sex→Diurnal | -0.331 | 0.645 | - | - |
| Sex→Pervasive | -0.144 | 0.852 | - | - |
| Age×sex→Diurnal | 0.011 | 0.867 | - | - |
| Age×sex→Pervasive | 0.021 | 0.761 | - | - |
| Home Conflict→ Diurnal | 0.127 | 0.178 | 0.028 | 0.686 |
| Home Conflict→ Pervasive | 0.174 | 0.068 | -0.039 | 0.507 |
| Home SES→ Diurnal | 0.051 | 0.629 | 0.007 | 0.844 |
| Home SES→ Pervasive | 0.029 | 0.824 | -0.017 | 0.757 |
| Home Adversity → Diurnal | -0.110 | 0.267 | 0.003 | 0.961 |
| Home Adversity → Pervasive | 0.054 | 0.646 | -0.084 | 0.308 |
| School-level Achievement→Diurnal | -0.160 | 0.083 | -0.278 | 0.052 |
| **School-level Achievement→Pervasive** | **-0.105** | **0.595** | **-0.356** | **0.003** |
| School Diversity→Diurnal | 0.032 | 0.816 | -0.013 | 0.844 |
| School Diversity→Pervasive | 0.147 | 0.359 | -0.066 | 0.321 |
| School Teacher Quality→Diurnal | -0.034 | 0.679 | | |
| School Teacher Quality→Pervasive | 0.159 | 0.111 | -0.110 | 0.228 |
| Neighborhood Poverty→Diurnal | 0.223 | 0.027 | -0.110 | 0.136 |
| Neighborhood Poverty→ Pervasive | -0.058 | 0.582 | -0.034 | 0.660 |
| Neighborhood Diversity→ Diurnal | -0.071 | 0.578 | -0.178 | 0.053 |
| Neighborhood Diversity→ Pervasive | 0.098 | 0.366 | -0.015 | 0.897 |
| Neighborhood (In)stability→ Diurnal | -0.037 | 0.764 | 0.023 | 0.737 |
| Neighborhood (In)stability→ Pervasive | -0.264 | 0.026 | -0.048 | 0.674 |

**Table S7.** Associations between sociecological context indicators and 'pervasive' and 'diurnal' cortisol in the younger portion of the sample (< 12 years of age, *n* = 135).

| Measures | *estimate* | $p_{uncorrected}$ | $p_{corrected}$ |
|---|---|---|---|
| Home Adversity → Diurnal | 0.050 | 0.675 | 0.9564 |
| Home Adversity →Pervasive | 0.033 | 0.793 | 0.9564 |
| Home Conflict→ Pervasive | 0.081 | 0.346 | 0.9427 |
| Home Conflict → Diurnal | -0.193 | 0.029 | 0.1740 |
| Home SES → Diurnal | -0.082 | 0.419 | 0.9427 |
| Home SES → Pervasive | 0.061 | 0.513 | 0.9564 |
| Neighborhood (In)stability → Diurnal | 0.018 | 0.917 | 0.9709 |
| Neighborhood (In)stability → Pervasive | -0.207 | 0.212 | 0.9427 |
| Neighborhood Diversity → Diurnal | -0.160 | 0.336 | 0.9427 |
| Neighborhood Diversity →Pervasive | 0.048 | 0.710 | 0.9564 |
| Neighborhood Poverty → Pervasive | -0.002 | 0.985 | 0.9850 |
| **Neighborhood Poverty → Diurnal** | **0.349** | **0.001** | **0.0090** |
| School Diversity → Diurnal | 0.036 | 0.797 | 0.9564 |
| School Diversity → Pervasive | 0.111 | 0.382 | 0.9427 |
| School Teacher Quality → Diurnal | -0.026 | 0.761 | 0.9564 |
| School Teacher Quality → Pervasive | 0.013 | 0.906 | 0.9709 |
| School-level Achievement → Diurnal | -0.039 | 0.680 | 0.9564 |
| **School-level Achievement → Pervasive** | **-0.564** | **0.001** | **0.0090** |

Note:→ = standardized regression paths; $p_{corrected}$ = *p* value corrected for multiple testing using the Bejamini-Hochberg false discovery rate (FDR) method

**Table S8.** Associations between sociecological context indicators * age and 'pervasive' and 'diurnal' cortisol.

| Measures | *estimate* | *p* |
|---|---|---|
| Home Conflict * age→ Diurnal | 0.017 | 0.626 |
| Home Conflict* age → Pervasive | 0.025 | 0.505 |
| Home SES* age → Diurnal | -0.002 | 0.970 |
| Home SES* age → Pervasive | -0.052 | 0.375 |
| Home Adversity* age → Diurnal | -0.058 | 0.183 |
| Home Adversity * age → Pervasive | -0.024 | 0.661 |
| School-level Achievement* age →Diurnal | 0.000 | 0.999 |
| School-level Achievement* age →Pervasive | 0.101 | 0.085 |
| School Diversity* age →Diurnal | -0.036 | 0.433 |
| School Diversity* age →Pervasive | 0.042 | 0.376 |
| School Teacher Quality* age →Diurnal | -0.034 | 0.679 |
| School Teacher Quality* age →Pervasive | -0.068 | 0.076 |
| Neighborhood Poverty* age →Diurnal | 0.028 | 0.586 |
| Neighborhood Poverty* age → Pervasive | -0.073 | 0.138 |
| Neighborhood Diversity* age → Diurnal | -0.071 | 0.275 |
| Neighborhood Diversity* age → Pervasive | 0.013 | 0.812 |
| Neighborhood (In)stability* age → Diurnal | 0.068 | 0.306 |
| Neighborhood (In)stability* age → Pervasive | -0.039 | 0.542 |

*Note:* the nine separate models were conducted each including the effects of one socioecological indicator, age and their product as predictors of diurnal and pervasive cortisol output; → standardized regression path.

**Table S9.** Associations between sociecological context indicators * puberty and 'pervasive' and 'diurnal' cortisol.

| Measures | *estimate* | *p* |
|---|---|---|
| Home Conflict * puberty→ Diurnal | 0.096 | 0.136 |
| Home Conflict* puberty → Pervasive | 0.010 | 0.853 |
| Home SES* puberty → Diurnal | -0.069 | 0.360 |
| Home SES* puberty → Pervasive | -0.056 | 0.491 |
| Home Adversity* puberty → Diurnal | -0.019 | 0.830 |
| Home Adversity * puberty → Pervasive | 0.013 | 0.878 |
| School-level Achievement* puberty →Diurnal | 0.070 | 0.341 |
| School-level Achievement* puberty →Pervasive | 0.060 | 0.619 |
| School Diversity* puberty →Diurnal | -0.055 | 0.461 |
| School Diversity* puberty →Pervasive | 0.024 | 0.667 |
| School Teacher Quality* puberty →Diurnal | 0.112 | 0.114 |
| School Teacher Quality* puberty →Pervasive | -0.101 | 0.249 |
| Neighborhood Poverty* puberty →Diurnal | -0.077 | 0.373 |
| Neighborhood Poverty* puberty → Pervasive | -0.102 | 0.209 |
| Neighborhood Diversity* puberty → Diurnal | -0.215 | 0.088 |
| Neighborhood Diversity* puberty → Pervasive | -0.020 | 0.859 |
| Neighborhood (In)stability* puberty → Diurnal | 0.202 | 0.069 |
| Neighborhood (In)stability* puberty → Pervasive | -0.108 | 0.311 |

Note: Measure of Pubertal Development: Participants rated their pubertal development on the Pubertal Development Scale [6]. Boys and girls rated their growth in height, growth in body hair, and skin changes, such as pimples, on a 4-point scale (1 = Not Yet Begun to 4 = Has Finished Changing). Males additionally rated deepening of voice and growth of facial hair, while girls rated breast development and if they had started menstruating (1 = No, 4 = Yes). A total pubertal status score was calculated by averaging across the items. Pubertal status spanned the full range (1–4) in both sexes, but was positively skewed. The pubertal status score was log transformed to correct for skew. Twenty-two participants were missing pubertal development scores due to incomplete survey response. On average, girls (M = 2.04, SD = 0.79) reported more advanced pubertal status than boys (M =1.82, SD = 0.62). 27% (*n* = 143 from 534 responses) of girls reported having begun menstruating; → = standardized regression path.

**Table S10.** Correlations between all CORT indicators and the nine socioecological context indicators.

| | W | AR | DS | PT | React | Recov | Hair | Sch A | Sch D | Sch T | Home A | Home SES | Home C | Neighb SES | Neighb D | Neighb S |
|---|---|---|---|---|---|---|---|---|---|---|---|---|---|---|---|---|
| W | 1.00 | | | | | | | | | | | | | | | |
| AR | -0.61 | 1.00 | | | | | | | | | | | | | | |
| DS | -0.31 | **0.56** | 1.00 | | | | | | | | | | | | | |
| P T | 0.25 | 0.07 | 0.22 | 1.00 | | | | | | | | | | | | |
| React | -0.10 | 0.03 | -0.12 | **-0.45** | 1.00 | | | | | | | | | | | |
| Recov | -0.02 | -0.06 | 0.14 | -0.01 | **-0.57** | 1.00 | | | | | | | | | | |
| Hair | 0.11 | 0.14 | **0.23** | 0.33 | -0.14 | 0.09 | 1.00 | | | | | | | | | |
| Sch A | 0.05 | -0.07 | **-0.24** | 0.07 | -0.04 | 0.13 | 0.03 | 1.00 | | | | | | | | |
| Sch D | 0.06 | 0.22 | 0.03 | 0.02 | -0.12 | 0.06 | 0.05 | -0.19 | 1.00 | | | | | | | |
| Sch T | -0.07 | 0.06 | 0.08 | 0.08 | -0.04 | 0.00 | 0.04 | 0.10 | **-0.26** | 1.00 | | | | | | |
| Home A | -0.05 | 0.15 | 0.03 | -0.02 | -0.07 | 0.11 | 0.07 | -0.06 | **0.31** | -0.09 | 1.00 | | | | | |
| Home SES | -0.04 | -0.16 | 0.20 | 0.05 | 0.02 | -0.05 | -0.05 | 0.08 | **-0.34** | 0.03 | **-0.55** | 1.00 | | | | |
| Home C | 0.07 | 0.14 | -0.02 | 0.13 | -0.14 | 0.12 | 0.02 | -0.07 | 0.06 | -0.09 | 0.06 | -0.08 | 1.00 | | | |
| Neighb SES | **0.26** | -0.27 | -0.22 | 0.00 | -0.12 | 0.02 | 0.04 | 0.08 | **-0.55** | 0.11 | **-0.37** | **0.47** | 0.06 | 1.00 | | |
| Neighb D | -0.05 | 0.10 | 0.13 | -0.04 | 0.03 | 0.08 | 0.04 | 0.17 | 0.21 | 0.00 | 0.04 | -0.04 | -0.02 | -0.16 | 1.00 | |
| Neighb S | -0.03 | 0.02 | -0.12 | -0.10 | 0.07 | -0.06 | -0.11 | 0.10 | -0.18 | 0.04 | -0.09 | 0.09 | -0.02 | 0.18 | **-0.30** | 1.00 |

**Note.** W = Waking Levels, AR = Awakening Response, DS = Diurnal Slope, Pre T = Pre-TSST levels, React = TSST Reactivity, Recov = TSST Recovery, Sch A. = School-level Achievement , Sch D = Diversity in school, Sch T = teacher characteristics, Home A = cumulative adversity, Home SES = parental socio-economic status, Home C = parental conflict, Neighb SES = neighborhood socio-economic status, Neigh D = neighborhood diversity, Neigh S = neighborhood stability, Underlined estimates = significant at the $p < .05$ level, **bold estimates** = significant at the $p < .01$ level, all estimates are calculated in the same model after accounting for age, sex, and age × sex.

**Table S11.** Final sample size after basic exclusions for diurnal, reactive and hair cortisol

| | | | | | |
|---|---|---|---|---|---|
| | **In-home diurnal cortisol** | | | | |
| | Day 1 | Day 2 | Day 3 | Day 4 | Day 5 |
| Sample 1 | 374 | 382 | 385 | 383 | 93 |
| Sample 2 | 344 | 353 | 354 | 343 | 74 |
| Sample 3 | 341 | 352 | 339 | 380 | 83 |
| | **In-lab reactive cortisol** | | | | |
| Sample 1 | 419 | | | | |
| Sample 2 | 410 | | | | |
| Sample 3 | 407 | | | | |
| Sample 4 | 408 | | | | |
| | **Hair Cortisol** | | | | |
| Sample 1 | 1188 | | | | |

**Table S12.** Descriptive statistics for log transformed diurnal, reactive and hair cortisol samples

| | *N* | *M* | *SD* | Skew | Kurtosis |
|---|---|---|---|---|---|
| In-lab sample 1 | 419 | .87 | .87 | 1.70 | 7.19 |
| In-lab sample 2 | 410 | 1.26 | .88 | .90 | 3.61 |
| In-lab sample 3 | 407 | 1.00 | .82 | 1.24 | 4.26 |
| In-lab sample 4 | 408 | .77 | .76 | 1.67 | 7.26 |
| At-home sample 1, day 1 | 374 | 1.85 | 086 | -1.00 | 4.85 |
| At-home sample 2, day 1 | 344 | 2.18 | .71 | .59 | 10.82 |
| At-home sample 3, day 1 | 341 | -.65 | .90 | 1.52 | 5.29 |
| At-home sample 1, day 2 | 382 | 1.82 | .83 | -1.12 | 5.09 |
| At-home sample 2, day 2 | 353 | 2.24 | .63 | -.79 | 2.35 |
| At-home sample 3, day 2 | 352 | -.60 | .91 | 1.10 | 2.06 |
| At-home sample 1, day 3 | 385 | 1.87 | .73 | -.85 | 3.15 |
| At-home sample 2, day 3 | 354 | 2.24 | .58 | .14 | .99 |
| At-home sample 3, day 3 | 339 | -.75 | .89 | .93 | 1.27 |
| At-home sample 1, day 4 | 383 | 1.86 | .74 | -1.51 | 4.87 |
| At-home sample 2, day 4 | 343 | 2.16 | .62 | -.63 | 3.10 |
| At-home sample 3, day 4 | 380 | -.43 | 1.17 | 1.13 | 1.17 |
| At-home sample 1, day 5 | 93 | 1.90 | .61 | .01 | -.06 |
| At-home sample 2, day 5 | 74 | 2.13 | .91 | 2.57 | 16.18 |
| At-home sample 3, day 5 | 83 | -.42 | 1.10 | 1.58 | 2.72 |
| Hair sample | 1188 | 1.20 | 1.23 | .77 | 1.50 |

*Note*. Cortisol values and *N*s correspond to data after exclusions were applied but before residualization for year of assay. Prior to log transformation, hair cortisol concentrations were measured in pg/ml and salivary cortisol concentrations were measured in nmol/L.

**Table S13.** Descriptive statistics for socio-ecological context measures.

| | School | | | Neighborhood | | | Home | | |
|---|---|---|---|---|---|---|---|---|---|
| | Achievement | Diversity | Teacher Quality | SES | (in)Stability | Diversity | Cumulative Adversity | SES (edu+income) | Parental Conflict |
| N | 1470 | 1470 | 1470 | 1470 | 1470 | 1470 | 1470 | 1470 | 1470 |
| min | 1 | 1 | 1 | 1 | 1 | 1 | 1 | 1 | 1 |
| max | 307 | 318 | 318 | 368 | 368 | 368 | 24 | 373 | 29 |
| range | 306 | 317 | 317 | 367 | 367 | 367 | 23 | 372 | 28 |
| median | 197 | 169.5 | 192 | 246 | 222 | 190 | 12 | 223 | 10 |
| mean | 186.6626 | 176.1170 | 185.5476 | 222.3939 | 214.1687 | 188.7667 | 12.4327 | 212.6592 | 11.9796 |
| SE | 2.5350 | 2.6305 | 2.6831 | 2.9690 | 2.9854 | 3.1403 | 0.2158 | 2.9642 | 0.2515 |
| SD | 97.1943 | 100.8536 | 102.8725 | 113.8350 | 114.4601 | 120.4001 | 8.2730 | 113.6496 | 9.6442 |

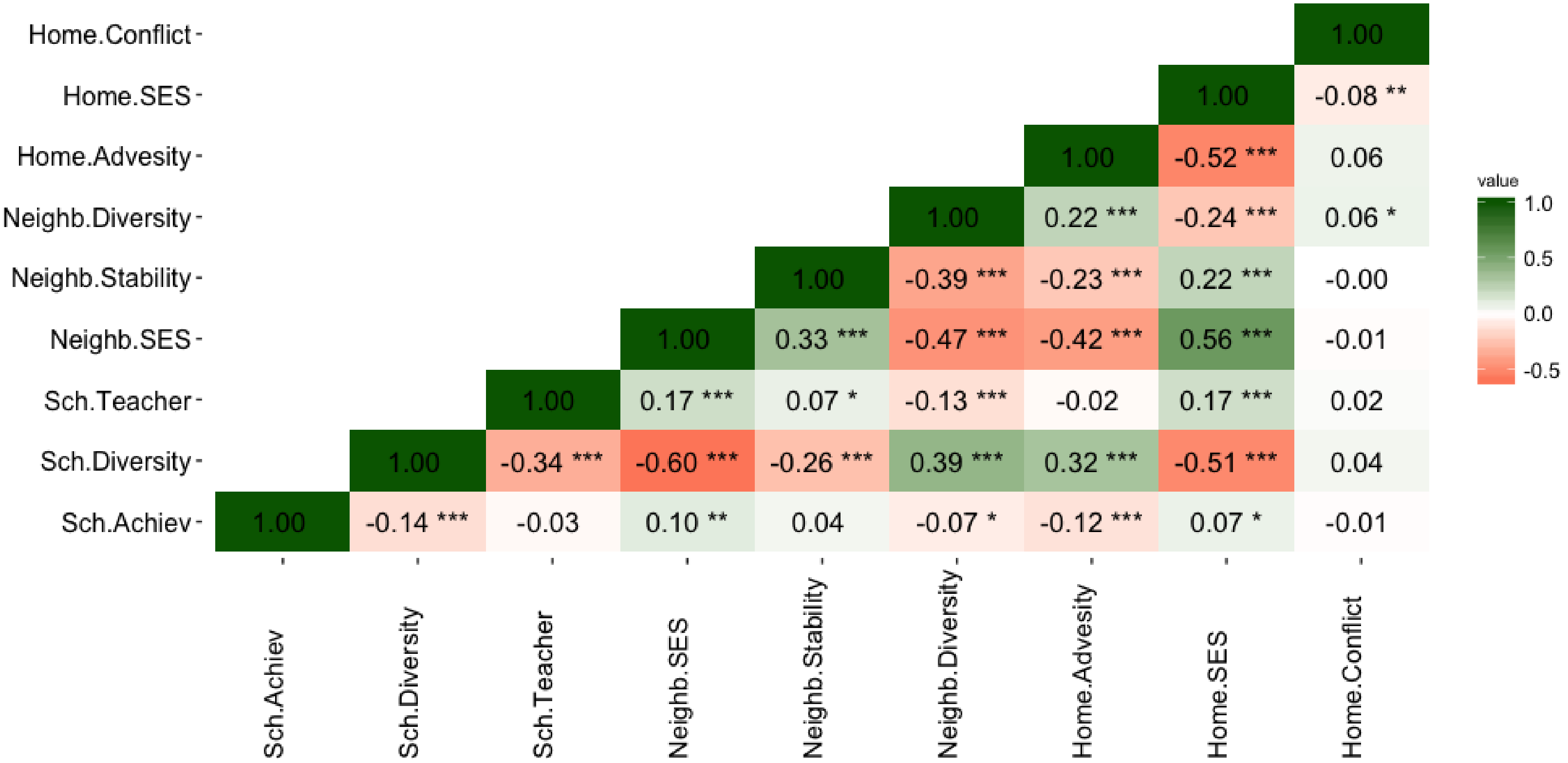

**Figure S1.** Correlations between the nine socioecological context indicators. Three indices reflected variation in the home environment: cumulative adversity (Home.adversity), parent socioeconomic status (Home.SES) and parental conflict (Home.conflict). Three indices covered several aspects of the school environment: school-level achievement (Sch.Achiev), school diversity (Sch.diversity) and teachers characteristics (Sch.teacher). Three indices reflected variation in numerous aspects of the neighborhood environment: neighborhood socioeconomic characteristics (Neighb.SES), residential stability (Neighb. Stability) and neighborhood diversity (Neighb.diversity); * = p< .05, ** = p< .01, *** = p< .001.

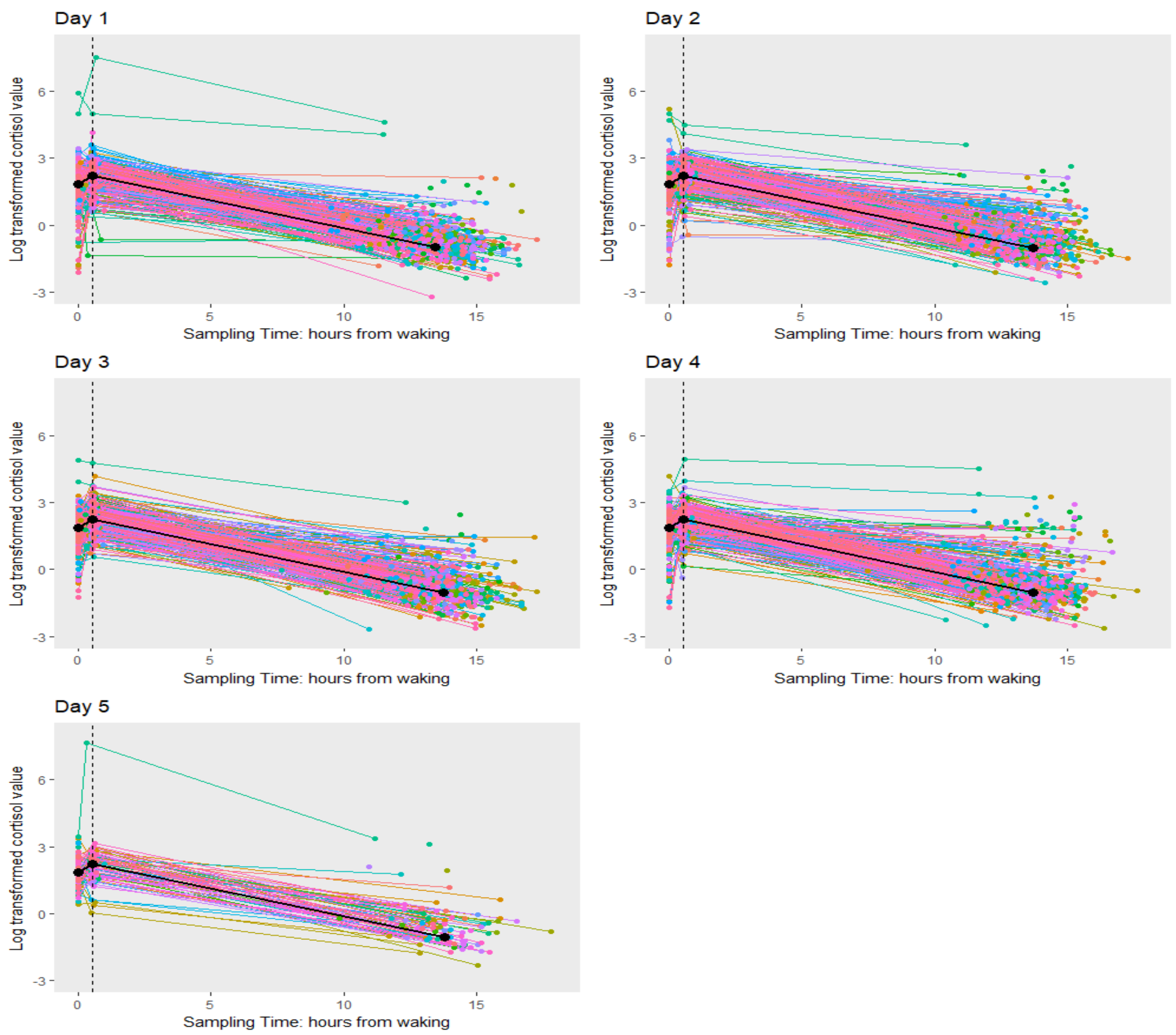

**Figure S2.** Intra and inter-individual variability in diurnal cortisol levels over the five sampling days (each colored line indicates a participant and each dot a sampling moment) and piecewise growth curves (black line) calculated across all sampling moments.

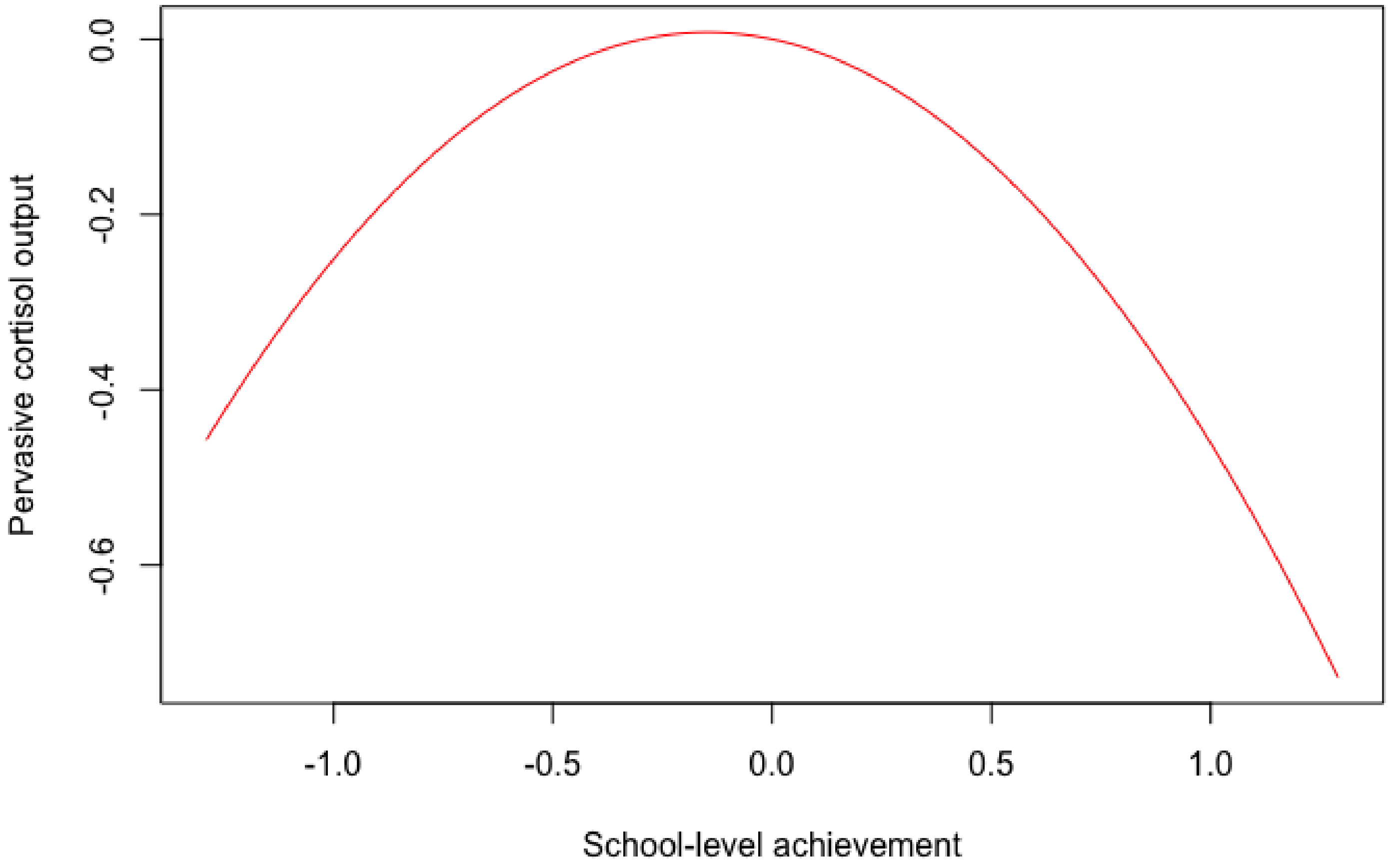

**Figure S3.** Regression line for the association between pervasive cortisol output and the quadratic term for school-level achievement

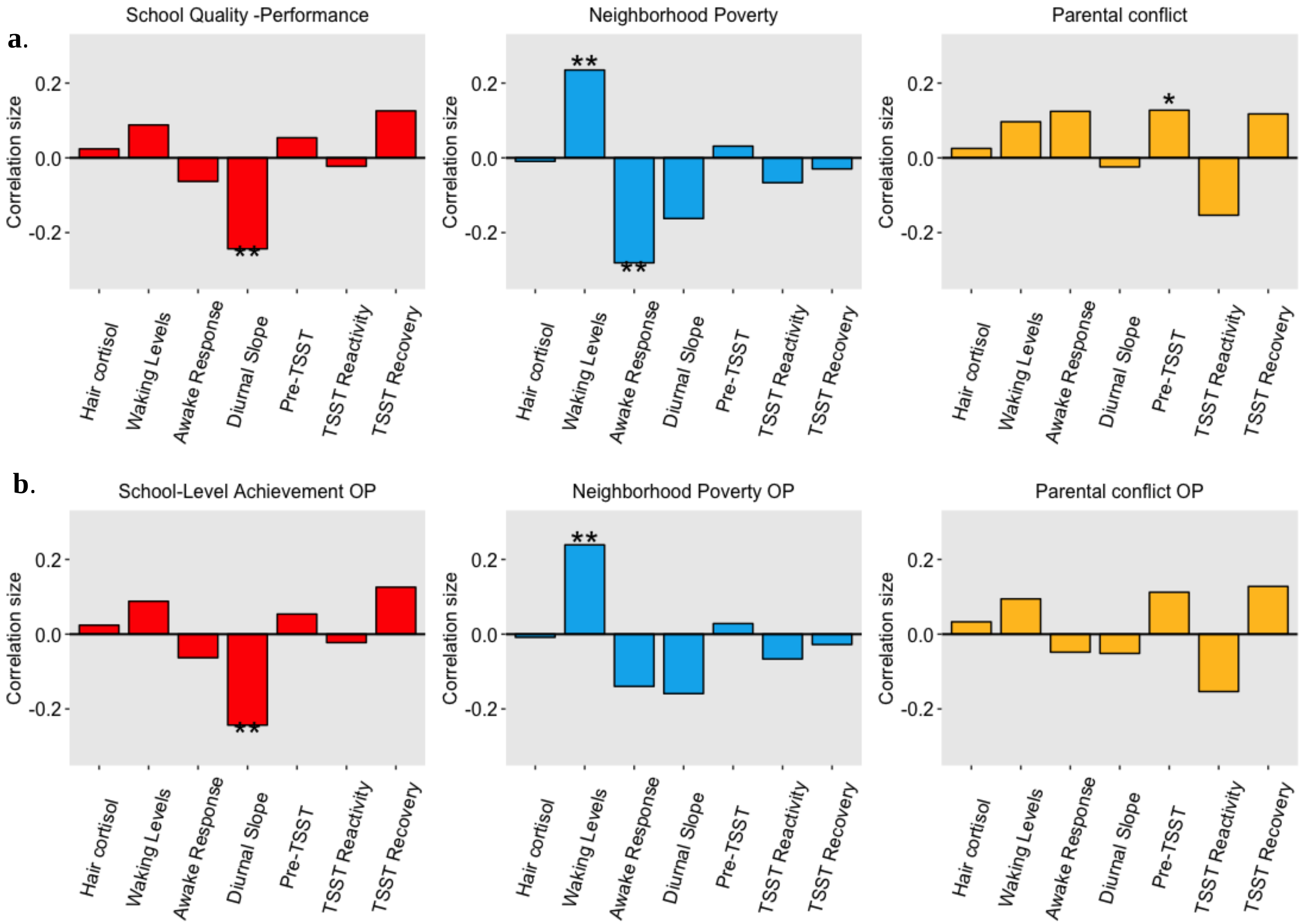

**Figure S4.** Associations between CORT latent factors and SES context measures in the final sample (panel a) and in the full sample, including participants showing off-phase CORT rhythm ($n$ = 36; panel b). Of the four associations shown in panel a, two were no longer observed when analyzing the full sample, rather than excluding due to being off-phase with respect to a naturally occurring circadian CORT rhythm. This was partly expected as the fact that the data points were off-phase with respect to the expected natural pattern of diurnal CORT fluctuation suggested that the samples had been collected later in the day, when levels of CORT had already started to decay, this hypothesis was corroborated by the fact that MEMs data was not available for this samples. Therefore, the lack of a significant association between awakening response and variation in neighborhood poverty potentially reflects a lack of compliance with the experimental procedure.

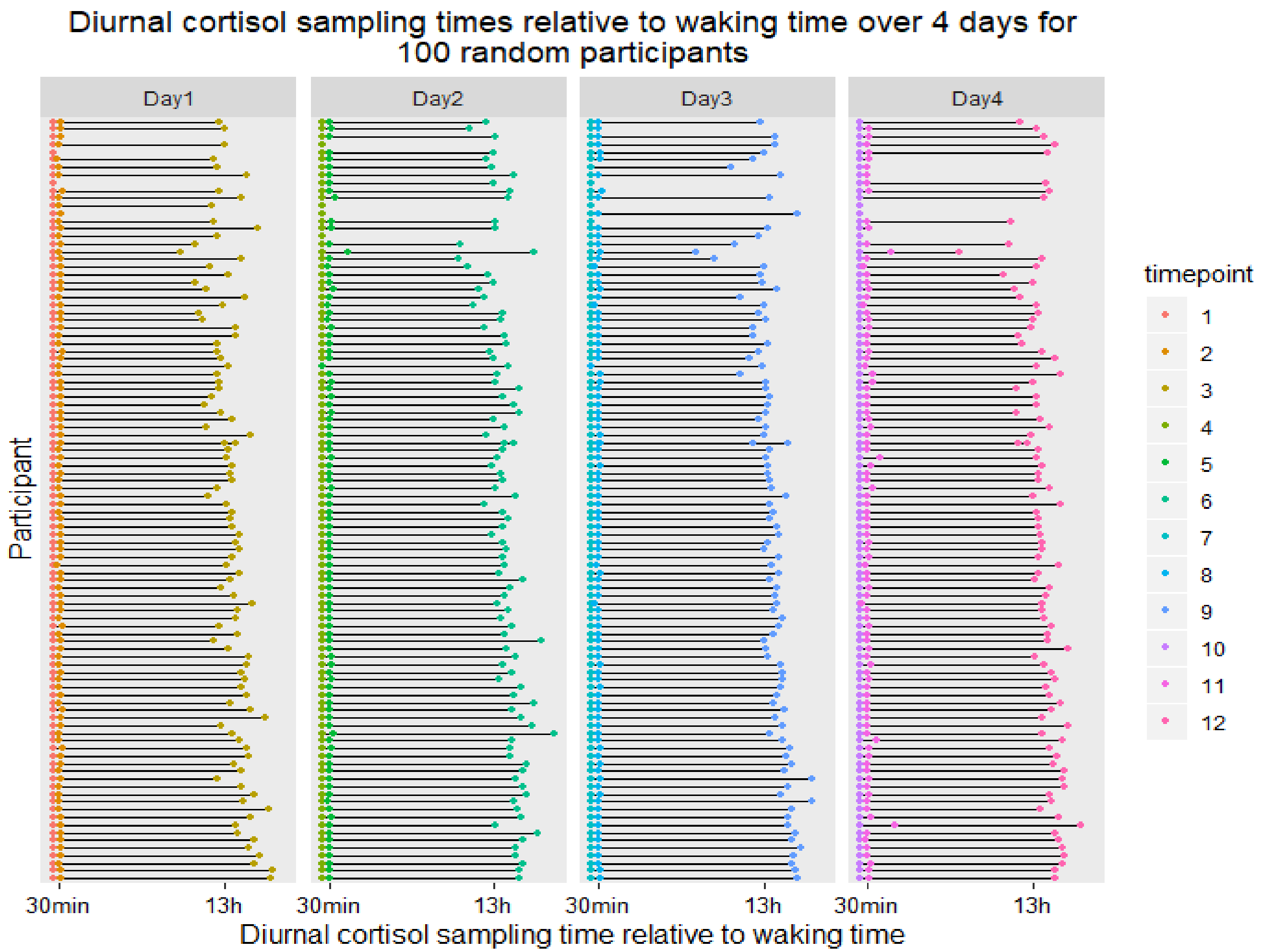

**Figure S5a.** Visual summary of data collection timeline of in-home diurnal cortisol for 100 randomly selected participants over the first four days (12 sampling moments)

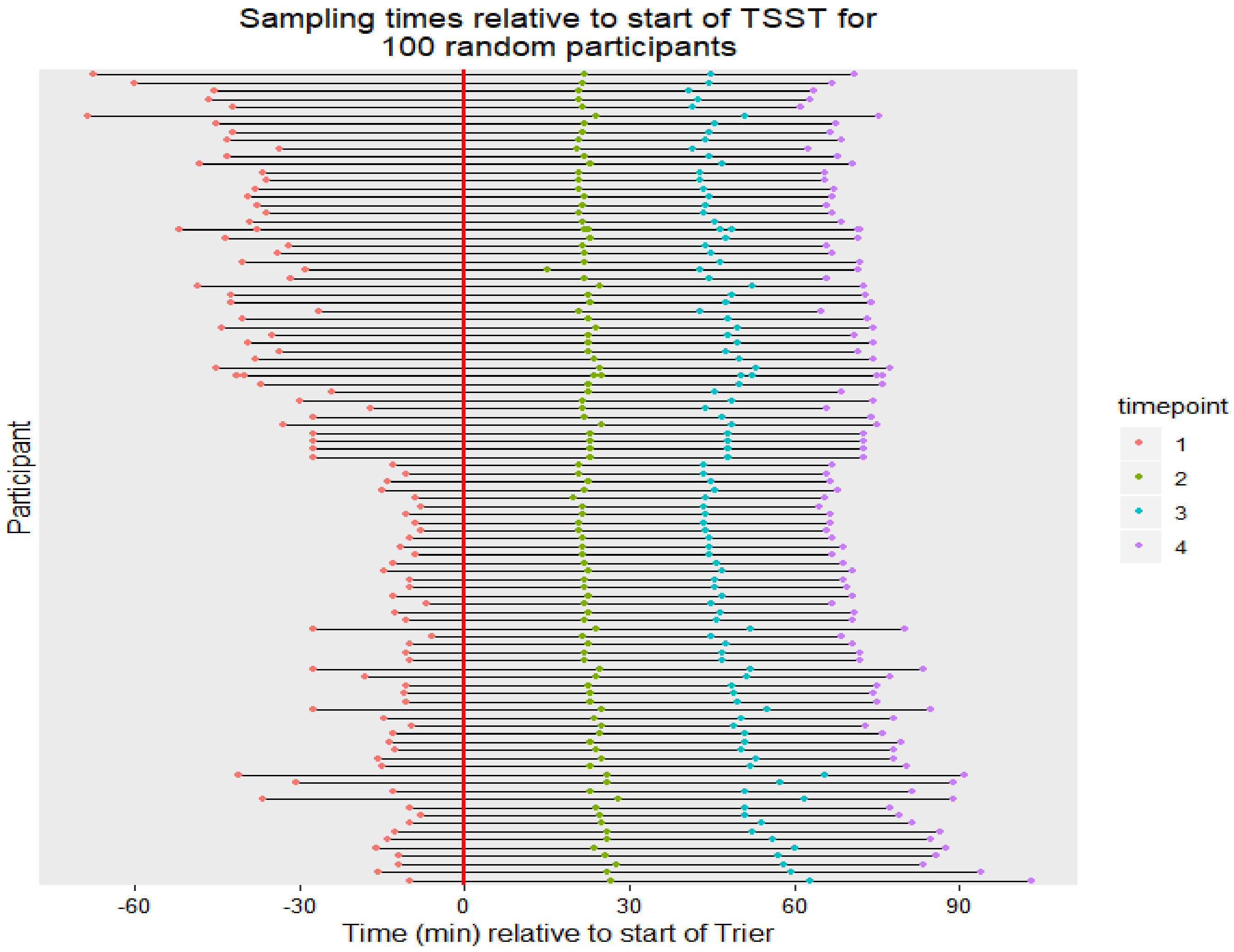

**Figure S5b.** Visual summary of data collection timeline of in-lab reactive cortisol for 100 randomly selected participants (4 sampling moments). The red line indicates the time of the Trier Social Stress Test (TSST).

**Figure S6.** Distributions of (**a**) log-transformed at-home diurnal cortisol samples, (**b**) log transformed in-lab reactive cortisol samples, (**c**) log transformed hair cortisol values, and **(d)** socio-ecological context measures.

**(a)** Distributions of log transformed at-home diurnal cortisol samples over the five sampling days.

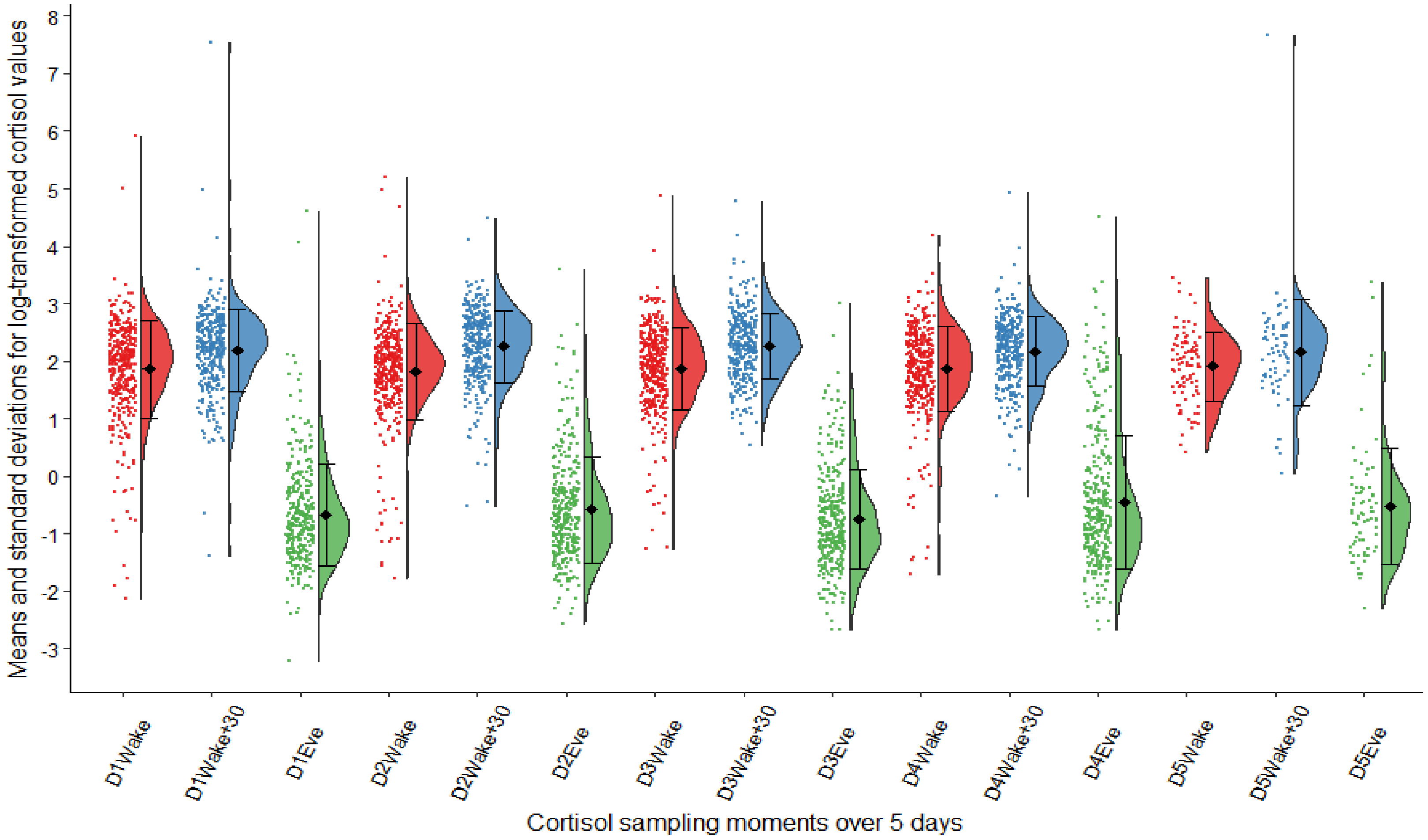

**(b)** Distributions of log transformed in-lab reactive cortisol samples.

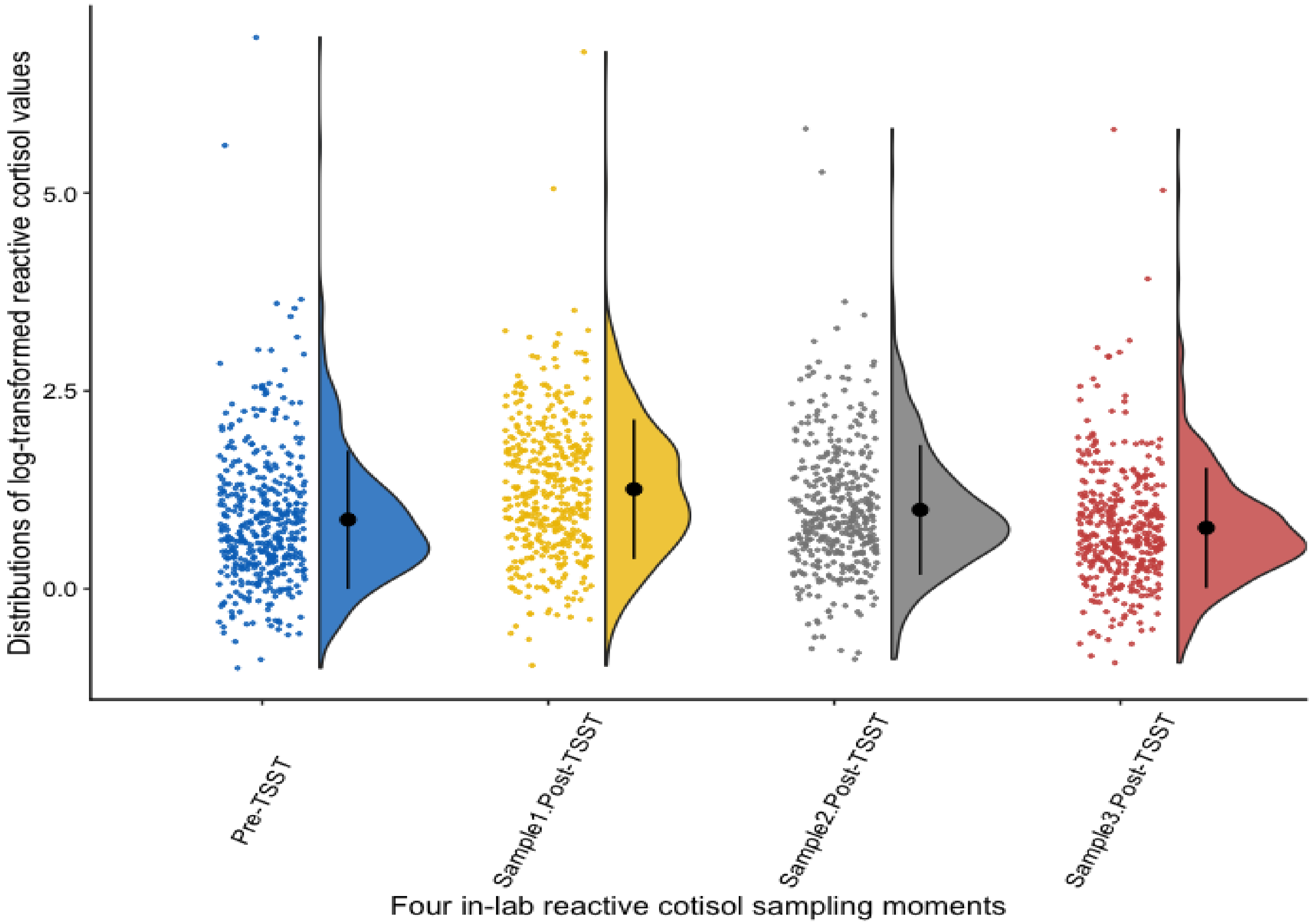

**(c)** Distribution of log transformed and batch-residualized hair cortisol.

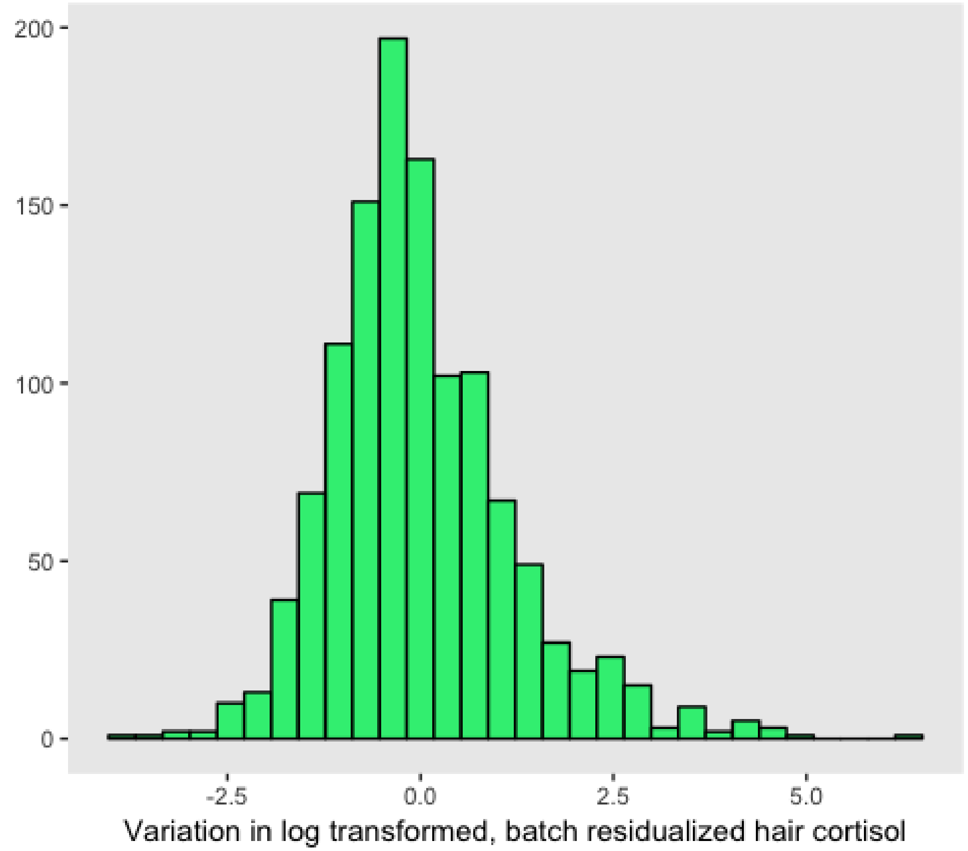

**(d)** Distributions of socio-ecological context measures

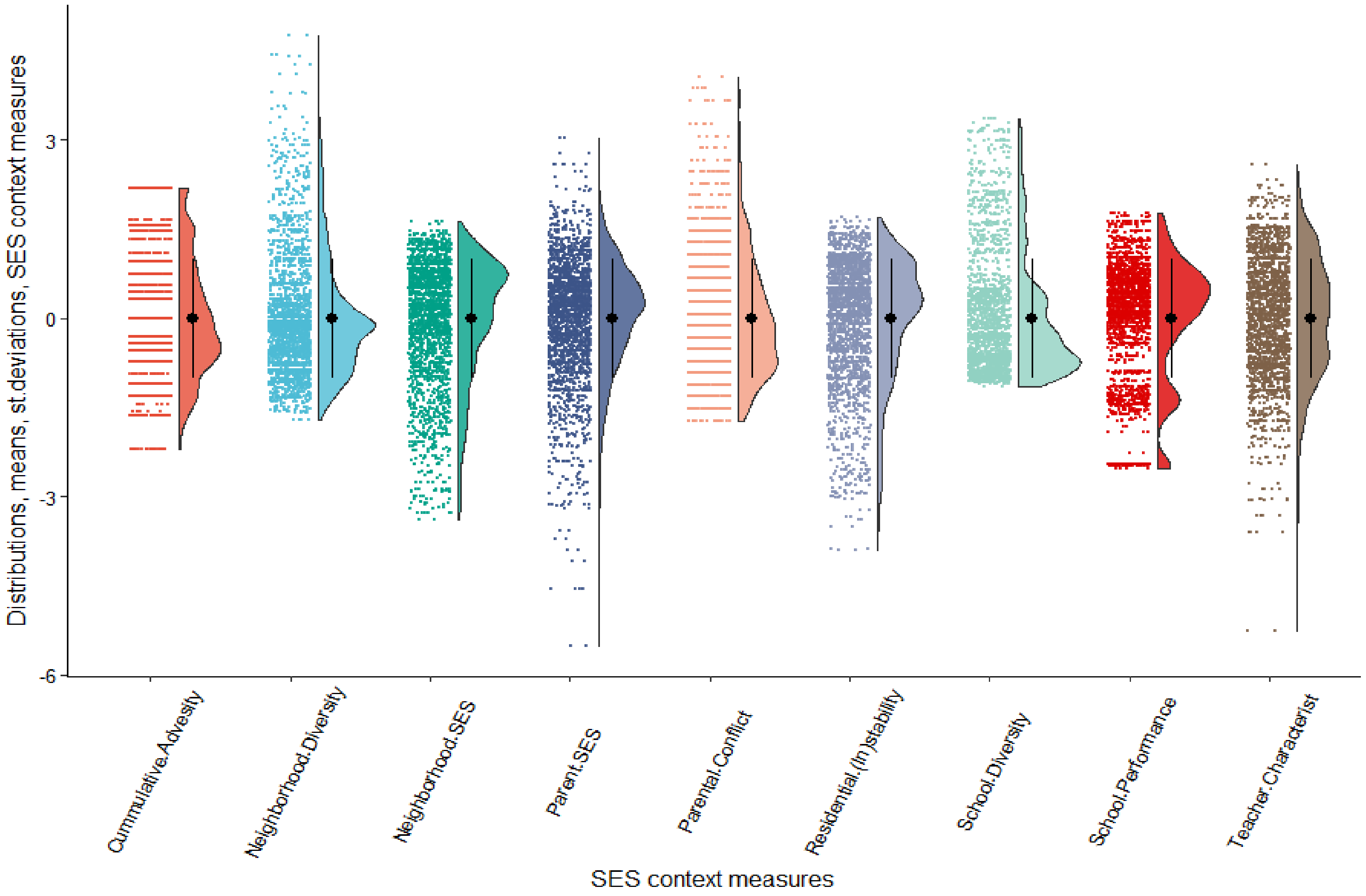

**Figure S7**. Model fit indices comparison for alternative latent growth models considering a range of turning points for (a) diurnal cortisol and (b) reactive cortisol trajectories.

(a) Diurnal

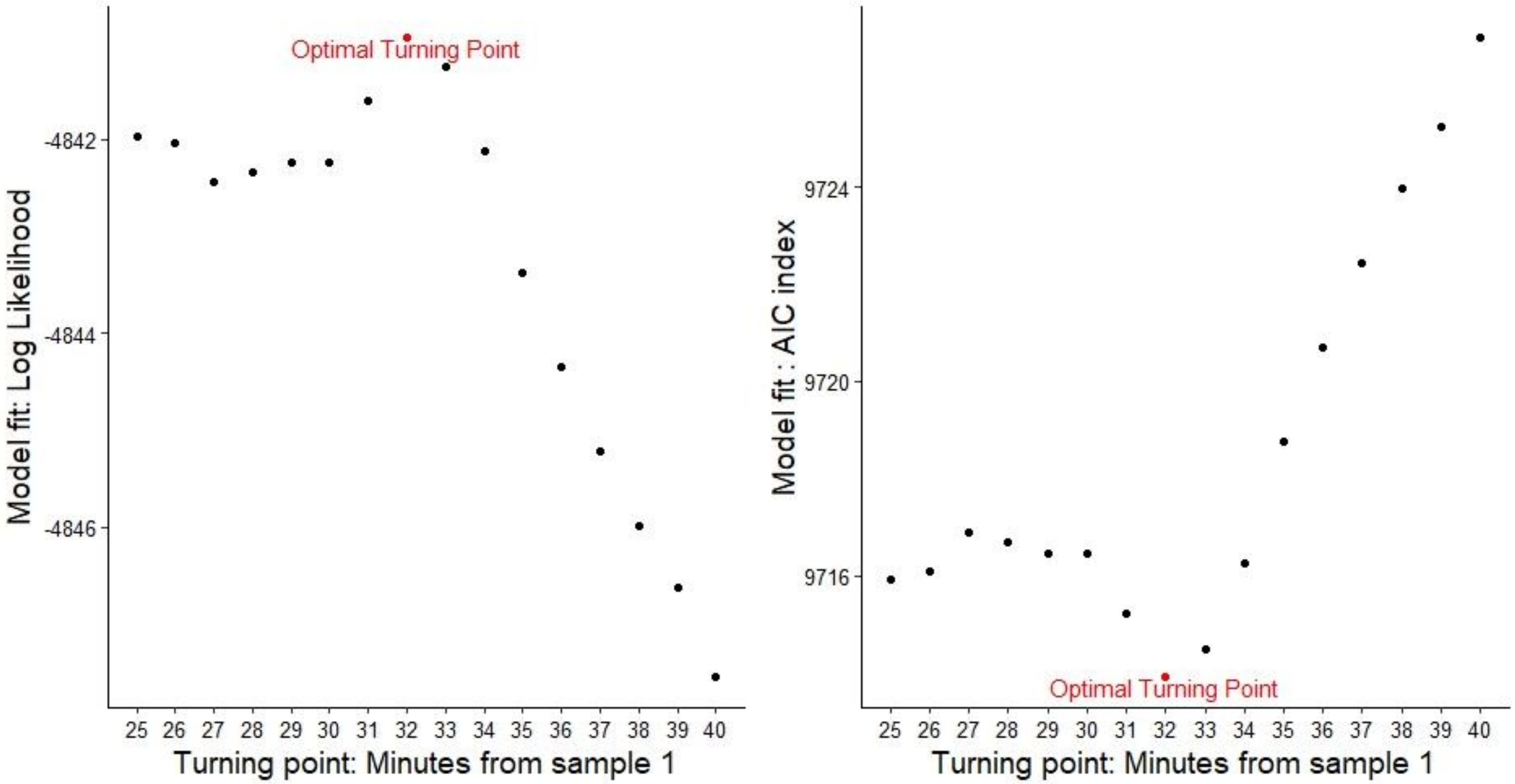

(b) Reactive

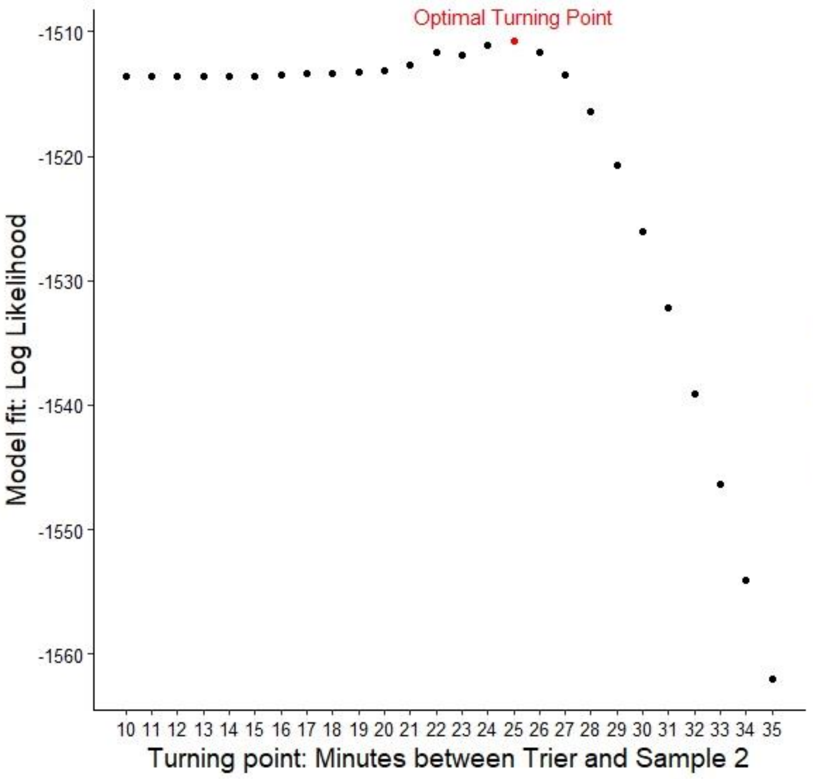

Optimal Turning Point
-1510
-1520
-1530
-1540
-1550
-1560
Model fit: Log Likelihood
10 11 12 13 14 15 16 17 18 19 20 21 22 23 24 25 26 27 28 29 30 31 32 33 34 35
Turning point: Minutes between Trier and Sample 2

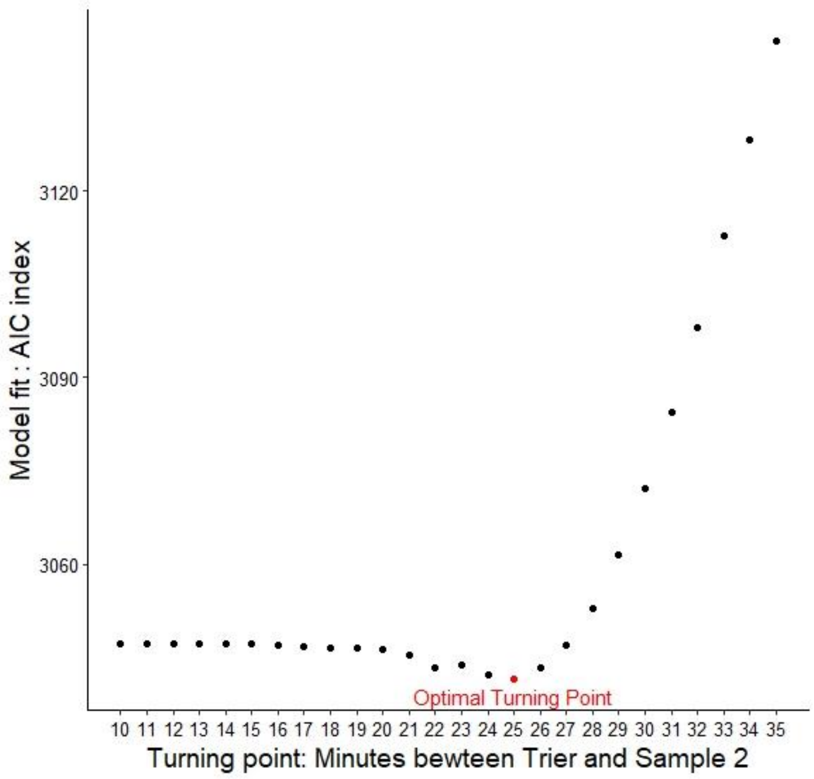

3120
3090
3060
Model fit : AIC index
Optimal Turning Point
10 11 12 13 14 15 16 17 18 19 20 21 22 23 24 25 26 27 28 29 30 31 32 33 34 35
Turning point: Minutes bewteen Trier and Sample 2

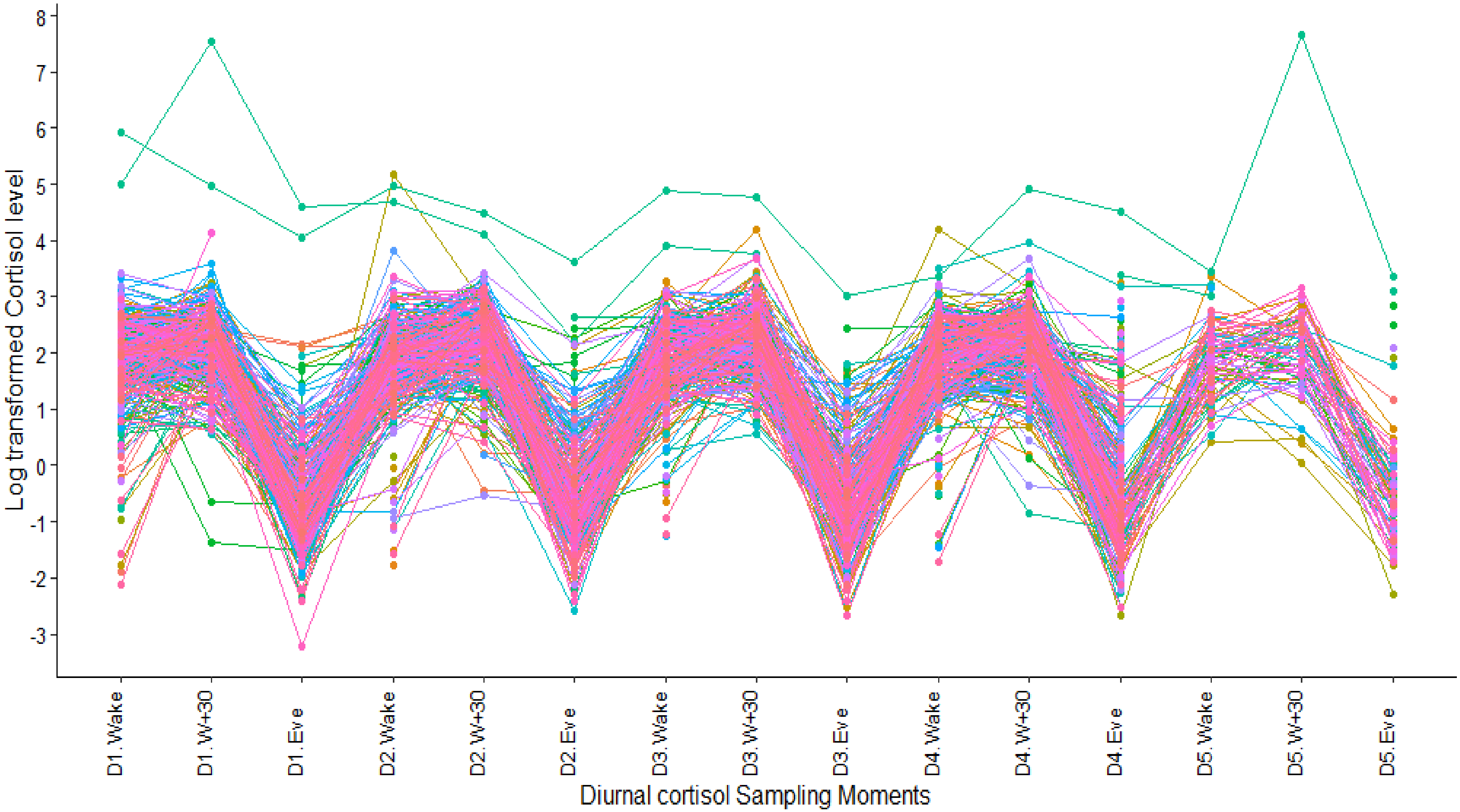

**Figure S8.a** Longitudinal coverage and level of log-transformed at-home cortisol samples after all exclusions over the 5-day collection period. Day1 = samples 1-3, Day 2 = samples 4-6, Day3 = samples 7-9, Day4 = samples 10-12, Day 5 = samples 13-15.

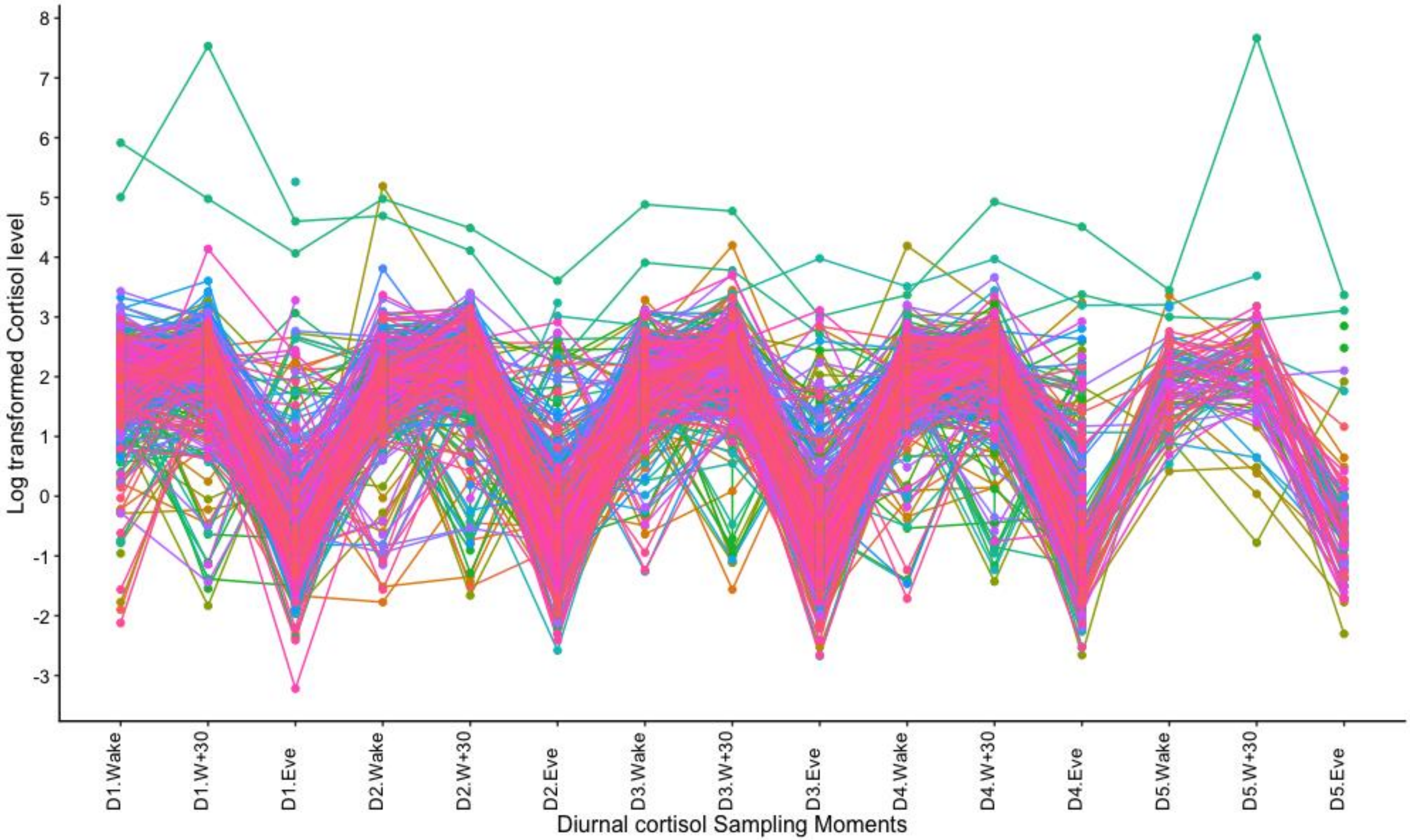

**Figure S8.b** Longitudinal coverage and level of log-transformed at-home cortisol **for all samples including those off-phase** over the 5-day collection period. Day1 = samples 1-3, Day 2 = samples 4-6, Day3 = samples 7-9, Day4 = samples 10-12, Day 5 = samples 13-15.
